# Long-read proteogenomics to connect disease-associated sQTLs to the protein isoform effectors of disease

**DOI:** 10.1101/2023.03.17.531557

**Authors:** Abdullah Abood, Larry D. Mesner, Erin D. Jeffery, Mayank Murali, Micah Lehe, Jamie Saquing, Charles R. Farber, Gloria M. Sheynkman

## Abstract

A major fraction of loci identified by genome-wide association studies (GWASs) lead to alterations in alternative splicing, but interpretation of how such alterations impact proteins is hindered by the technical limitations of short-read RNA-seq, which cannot directly link splicing events to full-length transcript or protein isoforms. Long-read RNA-seq represents a powerful tool to define and quantify transcript isoforms, and recently, infer protein isoform existence. Here we present a novel approach that integrates information from GWAS, splicing QTL (sQTL), and PacBio long-read RNA-seq in a disease-relevant model to infer the effects of sQTLs on the ultimate protein isoform products they encode. We demonstrate the utility of our approach using bone mineral density (BMD) GWAS data. We identified 1,863 sQTLs from the Genotype-Tissue Expression (GTEx) project in 732 protein-coding genes which colocalized with BMD associations (H_4_PP ≥ 0.75). We generated deep coverage PacBio long-read RNA-seq data (N=∼22 million full-length reads) on human osteoblasts, identifying 68,326 protein-coding isoforms, of which 17,375 (25%) were novel. By casting the colocalized sQTLs directly onto protein isoforms, we connected 809 sQTLs to 2,029 protein isoforms from 441 genes expressed in osteoblasts. Using these data, we created one of the first proteome-scale resources defining full-length isoforms impacted by colocalized sQTLs. Overall, we found that 74 sQTLs influenced isoforms likely impacted by nonsense mediated decay (NMD) and 190 that potentially resulted in the expression of new protein isoforms. Finally, we identified colocalizing sQTLs in *TPM2* for splice junctions between two mutually exclusive exons, and two different transcript termination sites, making it impossible to interpret without long-read RNA-seq data. siRNA mediated knockdown in osteoblasts showed two *TPM2* isoforms with opposing effects on mineralization. We expect our approach to be widely generalizable across diverse clinical traits and accelerate system-scale analyses of protein isoform activities modulated by GWAS loci.

## Introduction

Genome-wide association studies (GWASs) have identified thousands of associations influencing complex diseases ^1^; however, the main challenge limiting the use of GWAS data to uncover novel biology and new therapeutic targets is pinpointing causal genes. In recent years, it has become increasingly apparent that a substantial fraction of GWAS associations act by modulating alternative splicing (AS) ^2–4^. Genetic variants influencing AS are identified as splice quantitative trait loci (sQTLs) and several studies have linked sQTL to disease associations through colocalization approaches ^5–8^. However, it is not known to what extent the effects of GWAS loci are mediated by AS in general, and the identity of the downstream protein isoforms that mediate disease.

In a majority of functional genomics studies, splicing is characterized by algorithms such as LeafCutter ^9^ or rMATS ^10^, which quantifies local regions of splicing based on the relative abundance of introns (or exon-exon junctions) ^9^. These approaches have proven reliable as a way to globally quantify individual splicing events, and even in a reference annotation-free manner to discover novel splicing events. But, this information is only a partial picture—a majority of human genes express isoforms with multiple, distinct splicing events that can influence each other in cis, creating dependencies of splicing choices within the same transcript ^11,12^.

Unfortunately, short-read RNA-seq datasets can only return a probabilistic, not definitive, knowledge of isoform expression, many times with inaccuracies ^13^.

The influence of splicing in the genetics of complex disease is clear; however, it is often difficult to connect the events influenced by sQTLs to the full-length transcript isoforms they impact. This disconnect complicates efforts to interpret the effect of sQTLs and the design of experiments to test the role of specific isoforms for functional validation. Furthermore, the true impact of an sQTL on protein isoform function is unknown without direct measurement of the protein. It is only with knowledge of the number and sequence of protein isoform expression changes that one can determine how a GWAS locus mediates protein function from simple loss of protein stability to the generation of an alternative protein isoform with different functional activities, such as differential protein-protein interactions.

In recent years, long-read RNA-seq technologies have been shown to generate better predictions of proteoforms ^14^ including candidate novel protein isoforms ^15,16^. Here, we present a new approach that connects disease-associated sQTLs directly to the transcript and putative protein isoforms they impact. We accomplish this by integrating information from GWAS with large-scale sQTL datasets, which identifies hundreds of colocalized sQTLs, and directly cast the resultant sQTLs onto relevant isoform models derived from long-read RNA-seq data. In this process, sQTLs can be interpreted in terms of the disease-relevant protein isoforms that are correlated to a trait, enabling isoform-resolved studies from single-gene to systems scales. Such resolution facilitates hypothesis generation of individual or groups of isoforms playing a role in the clinical trait of interest.

As a proof of concept, we applied this sQTL-long-read contextualization approach to bone mineral density (BMD) GWAS ^17^ by leveraging sQTL from the Gene Tissue Expression (GTEx) project ^18^ and long-read proteogenomics in human fetal osteoblasts (hFOBs) ^19^ - a BMD relevant cell model. We identified 2,029 (643 novel) full-length protein isoforms from 441 protein-coding genes that are candidate effectors of BMD. To assess putative functions, we predicted complete open-reading frames and the effect of the associated protein isoforms on BMD. One of the genes identified through our approach was beta-tropomyosin (*TPM2*). Our analysis predicted that two different sets of isoforms characterized by the presence of two different mutually exclusive exons had opposing effects on mineralization, which we confirmed through isoform specific knockdown in hFOBs.

Our approach facilitates the interpretation of the effects of sQTLs to implicate isoforms likely involved in the regulation of BMD. This approach can be used for biomarker and novel therapeutic target identification, as well as understanding the splicing determinants of clinical traits, across the spectrum of human diseases.

## Results

An overview of our approach for systematic discovery of the transcript/protein isoforms potentially responsible for GWAS associations is shown in **Figure 1**. The approach uses a novel long-read proteogenomics approach ^15^ to increase the utility and interpretability of colocalized sQTLs.

**Figure 1:**
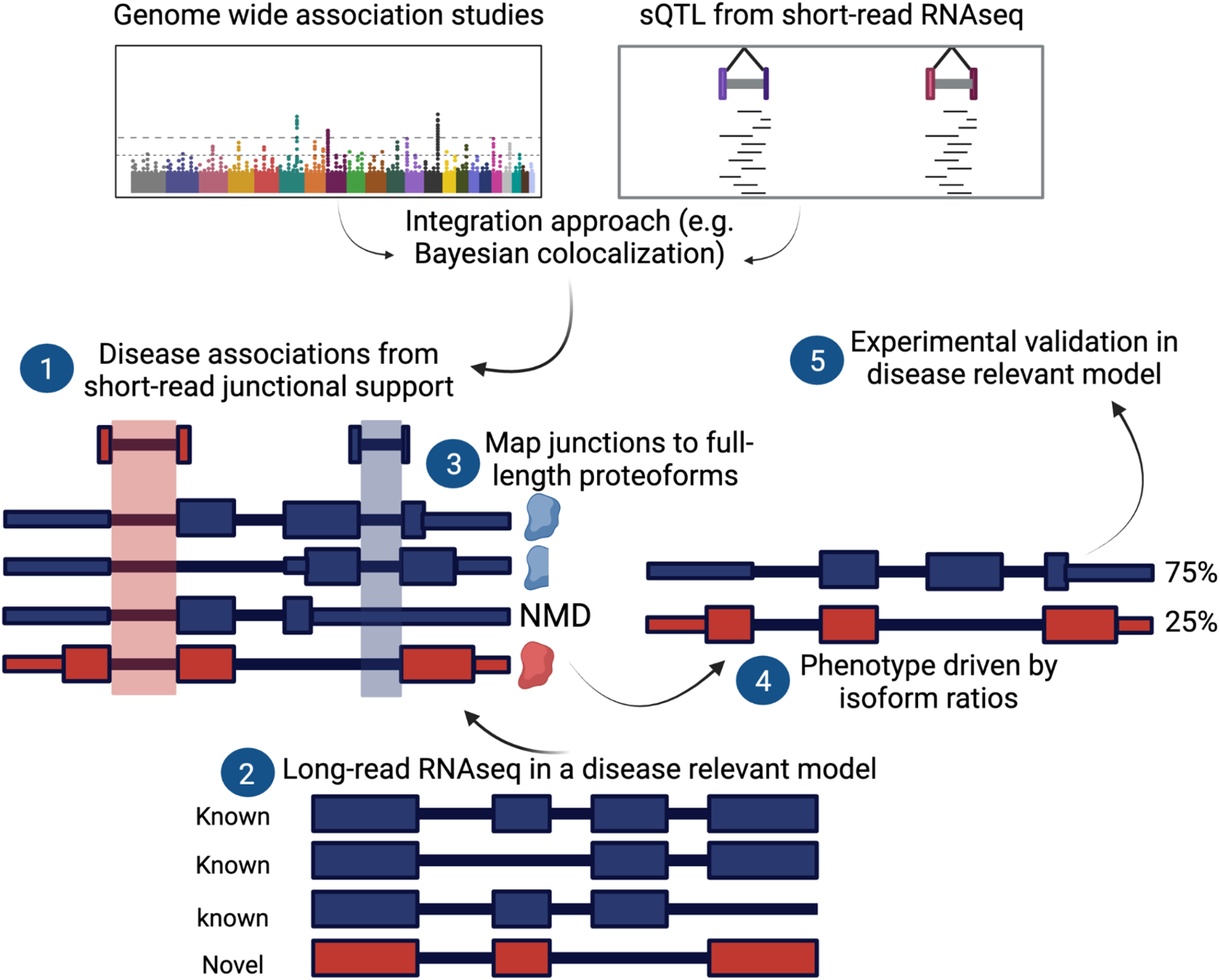
Overview of the approach to link genetically regulated splicing (sQTLs) to candidate protein isoform effectors. In step 1, disease associations were identified by integrating data from the latest BMD GWAS with sQTL data from 49 GTEx tissues using Bayesian colocalization analysis. In step 2, long-read RNA-seq data were generated from a disease relevant model identifying both known (blue) and novel (red) isoforms and their predicted open reading frames (ORFs), which in turn were used to map (step 3) the junctions identified in step 1 (red = novel, blue = known). Additionally, the impact of these junctions on ORFs was predicted (e.g truncation, nonsense mediated decay (NMD), or novel protein). Hypotheses generated from data in step 4 are then experimentally validated in the same disease model in step 5.

### Identification of genes potentially regulating BMD through splicing

To nominate splice events of potential relevance to BMD, we leveraged existing sQTLs across 49 tissues from the GTEx project ^5^. Population-scale transcriptomic datasets of bone or bone cells are scarce. However, prior studies have reported sharing of sQTLs across tissues, thus, we reasoned that a subset of the sQTL from GTEx would also be found in bone/bone cells and be of potential relevance to BMD ^3^. We performed Bayesian colocalization analysis using coloc ^20^, GTEx sQTLs, and the largest BMD GWAS performed to date, which identified 1103 independent associations ^17^ (**Supplemental note 1**). Overall, we found 732 protein-coding genes with colocalizing sQTLs (H_r_PP ≥ 0.75), denoted here as sGenes (**Figure 2A**). The colocalized sQTLs represent 1,863 distinct junctions with an average of 2.5 junctions per gene (**Figure 2A**). Over half of the sGenes (367, or 50%) have shared sQTLs across multiple tissues (**Table S1**). Examples of highly colocalizing sQTLs for *TCF7L2* (H_4_PP = 0.99) and *FHL3* (H_4_PP=0.99) are shown in **Figure 2A**. As illustrated in **Figures 2B and C**, when we mapped sQTLs to transcript annotations from GENECODE (v38), there were many transcripts potentially impacted, highlighting the challenge of interpreting the precise effects of sQTLs. Collectively, these results identify genes whose genetically regulated splicing alterations potentially mediate BMD associations.

**Figure 2:**
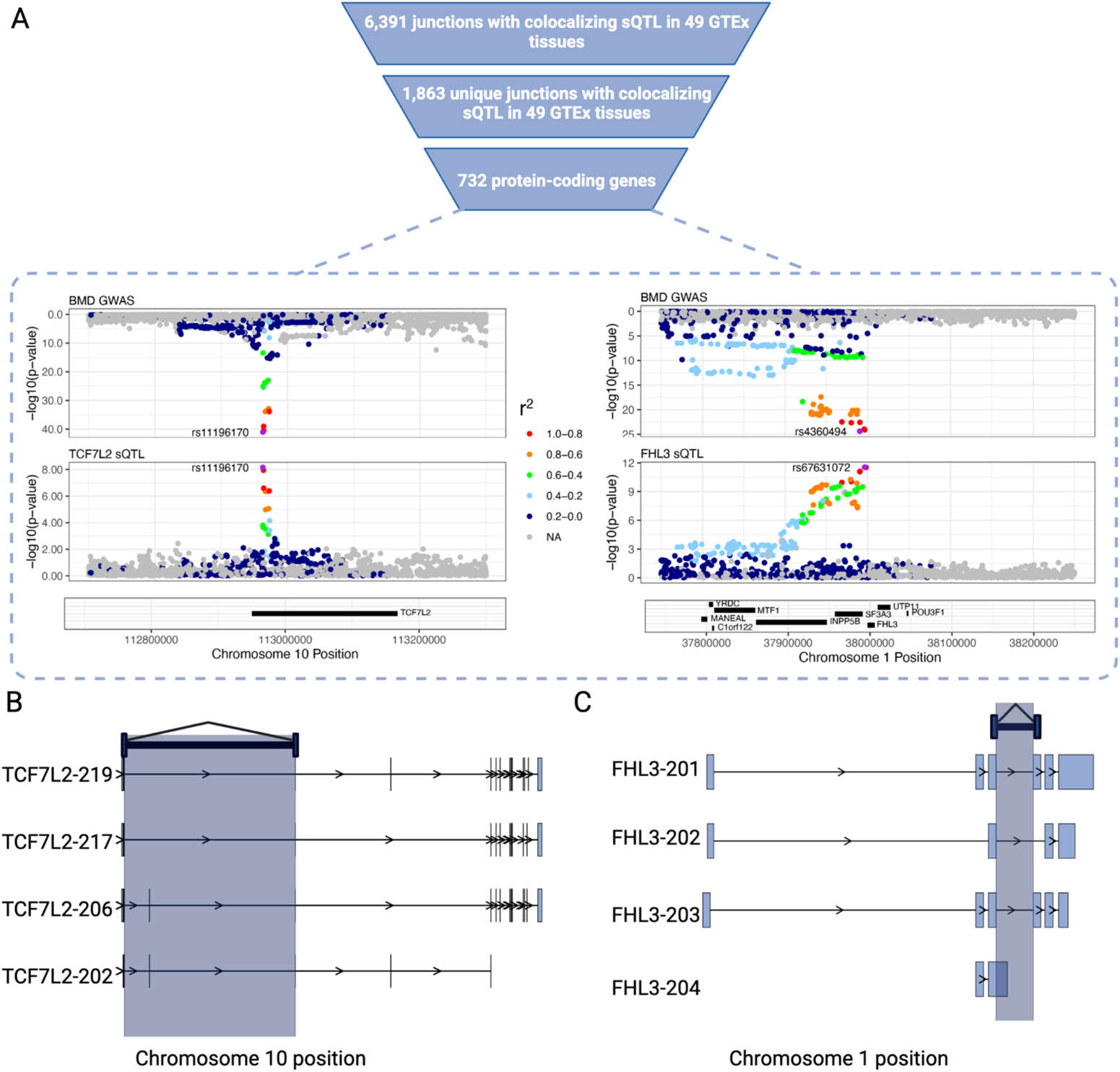
Identification of sQTLs colocalizing with BMD GWAS associations. A) Mirrorplots representing examples of highly 110 colocalizing sQTLs in TCF7L2 (H4PP = 0.99) and FHL3 (H4PP = 0.99). Mapping the junction with colocalizing sQTL to the referenctranscriptome (e.g., GENCODE) reveals multiple candidate isoforms in *TCF7L2* (B) and *FHL3* (C) that are potentially impacted by sQTLs.

### Characterizing the full-length transcriptome across osteoblast differentiation

In order to provide molecular context for the colocalized sQTLs, we generated an experimentally-derived and comprehensive map of isoform expression. Our goal was to enable the identification of the full-length transcript isoforms impacted by each colocalized sQTL and to ensure such isoforms were expressed in a cell-type relevant to BMD. To accomplish this, we generated deep coverage long-read RNA-seq data across osteoblast differentiation in hFOBs. Osteoblasts build bone and are critical regulators of bone development and maintenance ^21^. The hFOB cell line is a well characterized model of osteoblast differentiation capable of *in vitro* mineralization ^*22*^. Across 0, 2, 4, and 10 days of osteoblast differentiation, we generated a deep coverage full length transcriptome dataset (**Supplemental notes 2 & 4**), with collection of over 22 million high accuracy, full-length cDNA sequences using long-read RNA-seq on the PacBio Sequel II platform (**Figure S1A-B**) (see **Methods**). We detected 68,326 transcript isoforms from 12,068 genes. Transcripts of all lengths were evenly sampled (median: 2,074 nt, range: 87-8,787 nt). We found that 50,588 (74%) isoforms were known (annotated in GENCODE) and 17,738 (∼26%) were novel. Of the novel isoforms, 10,793 (61%) arose from new combinations of known splice sites (**Figure S1C**); whereas 6,580 (39%) arose from at least one novel splice donor or acceptor. Overall, this map of transcript isoforms in human osteoblasts is both comprehensive and critical for revealing cell-type-specific novel isoforms.

To characterize expression and splicing changes occuring during osteoblast differentiation, we used tappAS ^23^ and found that 2,034 genes (∼17% of all expressed genes) were differentially expressed (DE) and 3,539 (29% of all expressed genes) were differentially spliced, or undergoing differential isoform usage (DIU) (**Figure S1D**). Interestingly, DE and DIU genes were related to distinct GO terms. DE genes associated with bone-relevant processes such as positive regulation of bone mineralization (GO:0030501; FDR = 0.004), extracellular matrix organization (GO:0030198; FDR=8.88 × 10^−8^), and collagen-containing extracellular matrix (GO:0062023; FDR = 5.06 × 10^−11^) (**Figure S1D**). DIU genes, however, did not associate with annotated bone-relevant processes, but were enriched in terms related to the regulation of AS, including mRNA splicing and the spliceosome (GO:0000398; FDR = 0.0002) (**Figure S1D**). Together, these results suggest the presence of a splicing program acting independently of gene regulation during osteoblast differentiation, suggesting that genetic determinants of BMD could be acting through such splice-specific pathways.

### Connecting colocalized sQTLs to the transcript and protein isoforms they regulate through long-read derived isoform disease maps

Despite the wealth of global sQTL studies to date, few have connected sQTLs to the protein isoforms they impact. Here, we directly mapped colocalized sQTLs onto the protein isoform models generated by long read proteogenomics of a relevant disease model. We found that 836 junctions map exactly (shared splice donor/acceptor sites) to 2,349 isoforms (700 novel; ∼30%) in 459 protein-coding genes present in hFOBs (**Supplemental note 3**). Collectively, these sGenes are found within 362 of the 1,103 BMD GWAS associations (∼33% of the total), with 221 associations harboring one sGene and 141 harboring more than one sGene (**Table S2 and Supplemental note 5**). To our knowledge, this is the first global map of full-length isoform candidates that contribute to a human disease. Below, we highlight three examples demonstrating the power of long-read proteogenomics for interpretation of the effects of sQTLs.

### Colocalized sQTLs impacting novel isoforms

As we show above, it is now routine to discover hundreds of novel junctions corresponding to colocalized sQTLs. However, such novel sQTLs cannot be mapped to its source isoform(s) without experimental knowledge of full-length isoform expression, such as from long-read RNA-seq (**Figure 3A**). We detected 30 novel sQTL junctions, which were not found in the GENCODE reference, but mapped perfectly with one or more full-length novel isoforms detected in hFOBs. For example, we identified a novel junction with a colocalizing sQTL (H_4_PP = 0.99) in zinc finger protein 800 (*ZNF800*), a transcription factor expressed across a wide-range of cell-types that has been previously implicated in pancreatic beta cell development ^24^ and cardio-metabolic traits ^25^, but not in the regulation of BMD (**Figure 3B**). We mapped this junction to a novel transcript in hFOBs; however, it does not have a match to any isoform in the GENCODE database (v38) (**Figure 3C**). The lead variant (rs62621812) for both the BMD locus and sQTL is a rare missense variant (global minor allele frequency in 1000G = 0.005). Although rare, hFOBs were heterozygous for the variant and we observed that the novel *ZNF800* isoforms almost exclusively originated (21 of 22 long reads) from the haplotype harboring the alternative allele, and its expression decreases during hFOB differentiation (**Figure 3D**). The long *ZNF800* isoform containing the putative DNA binding domain is associated with increase in BMD, suggesting that its gene regulatory activities may be osteogenic.

**Figure 3:**
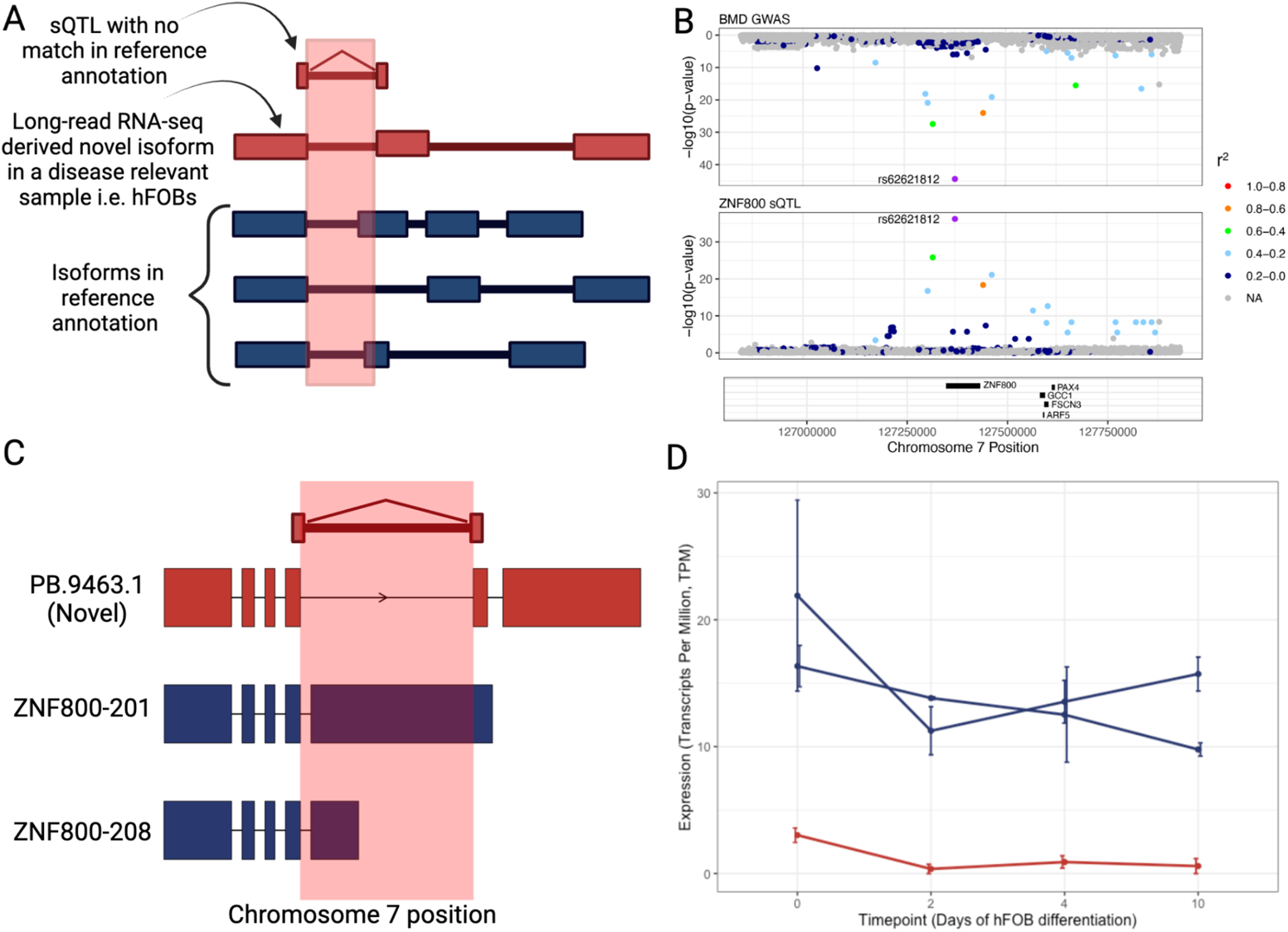
Colocalized sQTLs impact novel isoforms. A) Scenario in which a novel sQTL, having no match in isoforms within reference annotation (blue), mediaties its effect through a novel isoform (red) identified via long-read RNA-seq in hFOBs. B) Mirrorplot representing a highly colocalizing sQTL (H_4_PP = 0.99) in *ZNF800* where the most significant BMD GWAS SNP and the lead sQTL SNP are the same (rs62621812). C) Isoform models for *ZNF800* identified via long-read RNA-seq in hFOBs where the junction with colocalizing sQTL maps only to the novel isoform *PB.9463.1*. D) Expression of *ZNF800* isoforms across hFOB differentiation timepoints where red represents the novel isoform *PB.9463.1* and blue represents *ZNF800-201* and *ZNF800-208*. Bars represent standard error.

### Biological contextualization of isoforms corresponding to known sQTLs

Ostensibly, there is a clear path to contextualize sQTLs containing annotated junctions, as these junctions have a direct mapping to at least one isoform in the reference annotation. However, a single sQTL could map to multiple annotated isoforms and it is not known if all isoforms or only a subset is relevant in mediating the trait of interest, a common case of isoform ambiguity in sQTL datasets (**Figure 4A**). We found a total of 614 sQTLs (73% of 836) mapping to multiple annotated isoforms in the GENCODE transcriptome (v38). To hone in on the most relevant isoforms for the trait of interest, we leveraged information about which isoforms were expressed in the hFOB transcriptome map.

**Figure 4:**
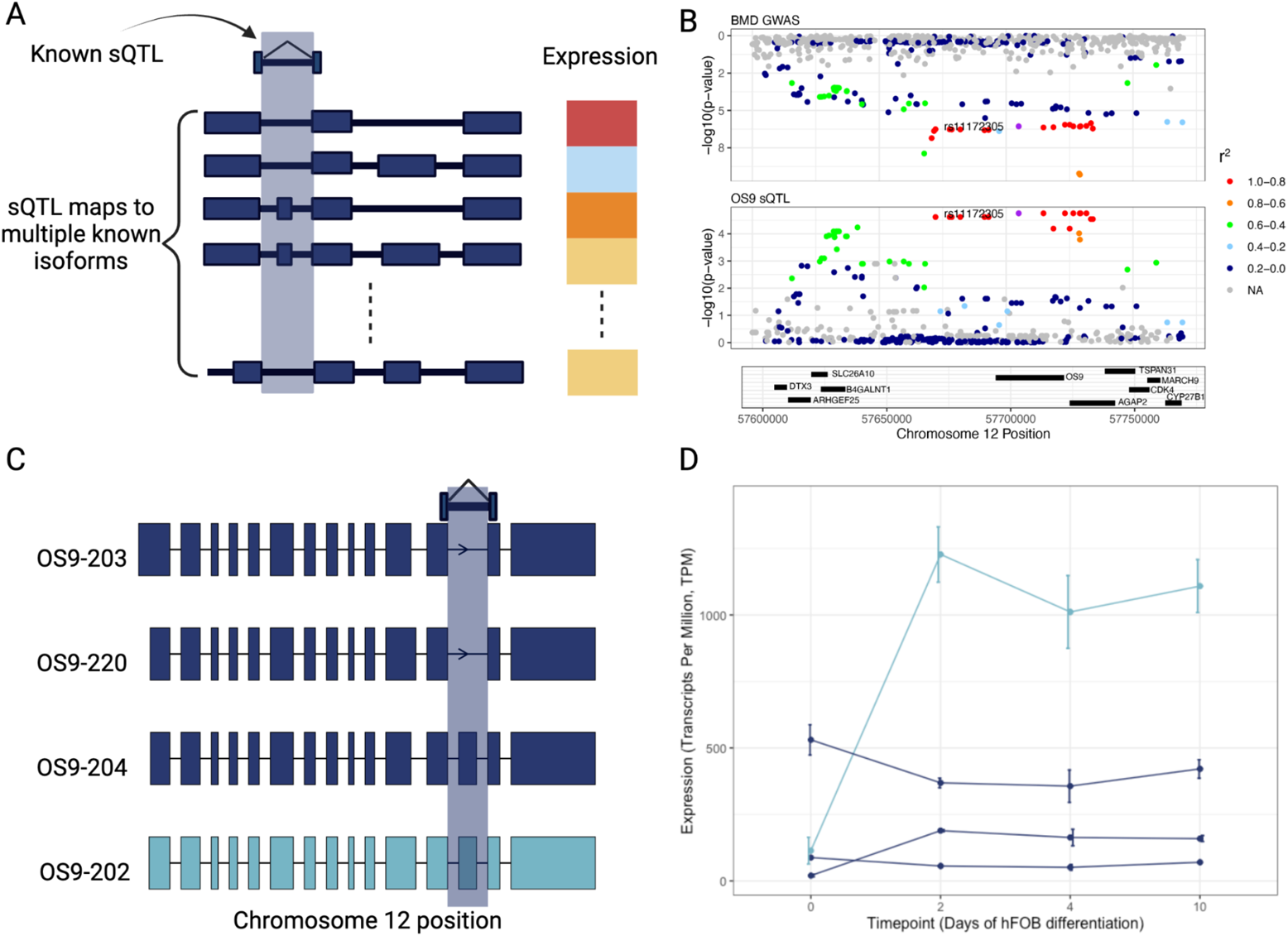
Contextualization of isoforms corresponding to known sQTLs. A) Scenario in which a known sQTL (match in isoforms within reference annotation) shown in blue map to multiple annotated isoforms (blue) and it is not known if all isoforms or only a subset may be relevant in mediating the trait of interest. Therefore, full-length expression can be leveraged (scaled from red (high expression) to blue (low expression)). B) Mirrorplot representing a colocalizing sQTL (H_4_PP = 0.86) in *OS9*. C) Isoform models for *OS9* identified via long-read RNA-seq in hFOBs where the junction with colocalizing sQTL maps to two isoforms *OS9-220* and *OS9-203*. D) Expression of *OS9* isoforms across hFOB differentiation timepoints where light blue represents isoform *OS9-202* and blue represents the rest of the isoforms. This example highlights that putting these sQTLs within a biological context can nominate more directly potential causal isoforms Bars represent standard error.

For example, the sQTL-to-isoform mapping of amplified in osteosarcoma 9 (*OS9*) illustrates the power of providing full-length isoform context. *OS9* has not been directly implicated in the regulation of BMD, but we observed a colocalizing sQTL (H_4_PP = 0.86) (**Figure 4B**) that corresponds to the skipping of exon 13. This sQTL maps to 24 isoforms in the GENCODE reference annotation, leaving open the question of which isoform(s) may be specifically relevant to bone cells. Within the hFOB context, we found four *OS9* isoforms expressed in hFOBs: *OS9-203* and *OS9-220*, which both exclude exon 13, and *OS9-202* and *OS9-204*, which include exon 13 (**Figure 4C**). We found that *OS9-220* and *OS9-202* are the dominantly expressed isoforms and exhibit an isoform switch during osteoblast differentiation with the exon 13-included form, *OS9-202*, dramatically increasing in abundance from day 0 to 2, relative to *OS9-220*, suggesting a direct role in driving osteoblast differentiation (**Figure 4D**). Interestingly, independent evidence from the International Mouse Phenotyping Consortium (IMPC) ^26^ showed that *Os9* knockout mice show significant changes in skeletal phenotypes including abnormal cranium morphology (p = 1.40 × 10^−5^; both sexes), abnormal tooth morphology (p = 9.12 × 10^−5^; both sexes), and vertebral fusions (p = 3.92 × 10^−5^; males only).

### Known sQTLs impacting novel isoforms

Disease-relevant isoform expression information can narrow the possibilities of isoforms within which the annotated junctions of sQTLs map. However, just as importantly, annotated junctions of colocalized sQTLs can be found to map to novel isoforms, meaning that the local splicing event is known, but the associated full-length protein isoform to which it is derived is novel (**Figure 5A**). These “indirectly” novel sQTLs may be an underappreciated source of novel isoforms regulated by GWAS loci. We found that a total of 383 (46%) colocalized sQTLs were found in known isoforms only, but interestingly 350 (42%) can map to both known and novel isoforms. Strikingly, we found that 103 (12%) were exclusively mapping to, and thus, potentially explained by novel isoforms.

**Figure 5:**
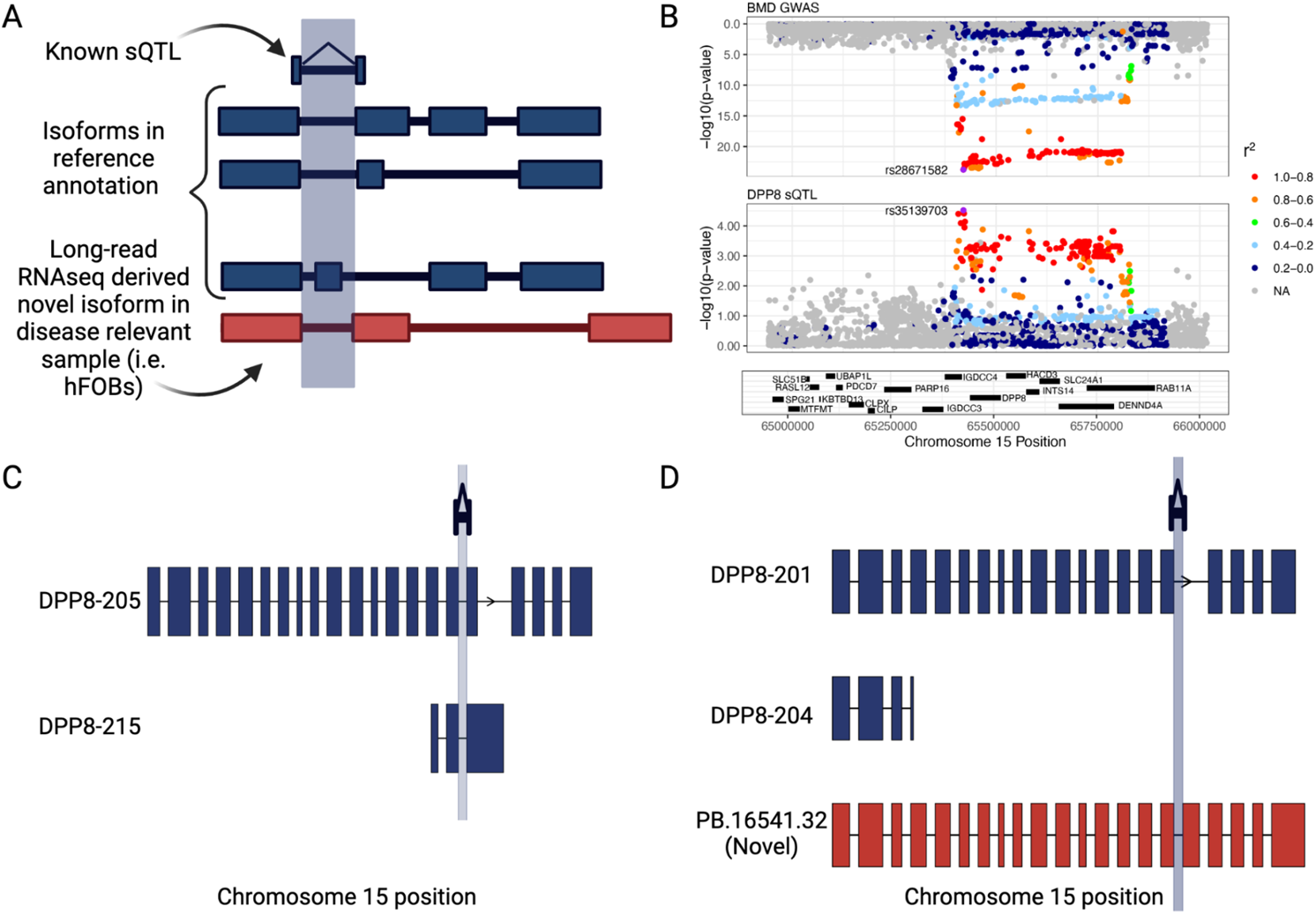
Known sQTLs impact novel isoforms in the biological context. A) Scenario in which annotated junctions of colocalized sQTLs can actually be found to map to novel isoforms, meaning that the local splicing event is known (blue), but the associated full-length protein isoform to which it is derived is novel (red). B) Mirrorplot representing a colocalizing sQTL (H_4_PP = 0.83) in *DPP8*. C) Isoform models for *DPP8* in GENCODE v38 containing the junction (blue). D) Isoform models for *DPP8* identified via long-read RNA-seq in hFOBs where the junction with colocalizing sQTL maps exclusively to a novel isoform *PB.16541.32*.

Since these known sQTLs map to novel isoforms, they might be “re-annotated” as candidate novel protein isoforms. An example includes dipeptidyl peptidase 8 (*DPP8)*, for which we found a colocalizing sQTL (H_4_PP = 0.83) corresponding to inclusion of exon 17 (**Figure 5B**). The associated junction can be mapped in GENCODE isoforms *DPP8-205* and *DPP8-215* (**Figure 5C**), however, neither are expressed in hFOBs (**Figure 5D**). Rather, the junction maps to a novel isoform (*PB.16541.32)* (**Figure 5D**). The alternative isoform of the sQTL corresponds to skipping of exon 17 and is found in isoform *DPP8-201*, which is associated with a decrease in BMD. Interestingly, in data from the IMPC ^26^, *Dpp8* knockout mice have decreased BMD.

### Enrichment of splicing functions among BMD sQTLs and presence of a putative splicing regulatory network

Among all 459 genes with a colocalizing sQTL, we found an enrichment in alternative splicing (UniProt Keyword KW-0025 6.39 × 10^−17^, RNA splicing). Splice factors (SFs) regulate splicing in trans and have been found to comprise 99 genes to date ^27^ (**Figure 6A**). Of the 99 annotated SFs, 91 were expressed in hFOBs. We found 5 splice factors that show a colocalizing sQTL within our long-read hFOB data: *MBNL1, PCBP2, DDX5, PTBP1*, and *HNRNPM*. Three of these (60%) show differential expression across hFOB differentiation: *DDX5, PTBP1*, and *HNRNPM* and one showed differential isoform usage: *DDX5* (**Supplemental note 6**).

**Figure 6:**
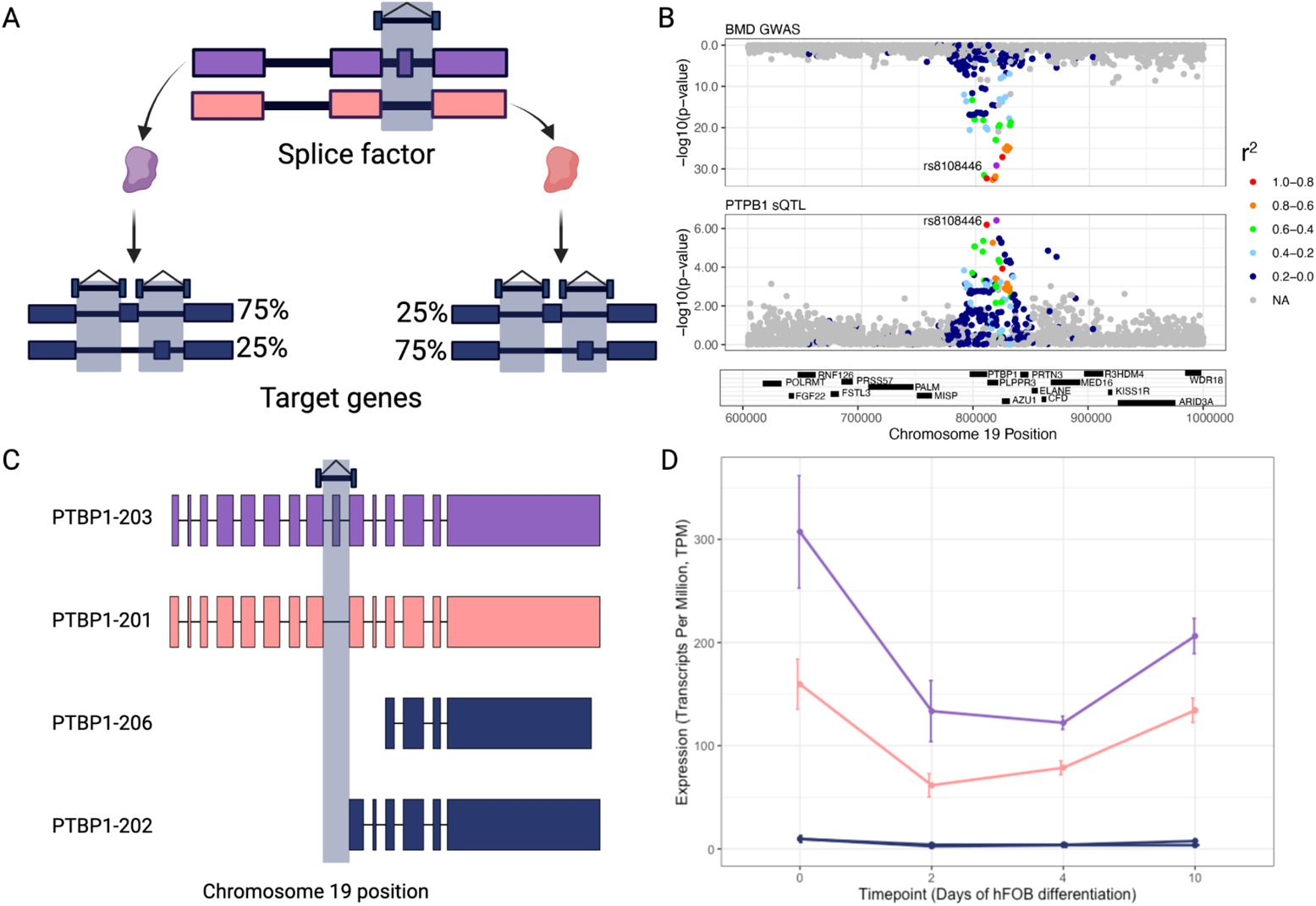
Enrichment of lead sQTLs in splice factor binding sites. A) Splice factors are regulated by sQTLs and in turn regulate the splicing of target genes (that can also contain sQTLs). B) Mirrorplot showing a significantly colocalizing sQTL adjacent to exon 9 in the splice factor *PTBP1*. C) Isoform models of *PTBP1* with the junction coordinates highlighted. D) Expression of all *PTBP1* isoforms including *PTBP1-203* (purple) and *PTBP1-201* (pink). Bars represent standard error.

One example of a putative splicing regulator potentially involved in the regulation of BMD is polypyrimidine tract binding protein (*PTBP1*), which has been extensively studied as a global repressor of exons ^28^ and is implicated in complex diseases such as atherosclerosis ^29^. In our colocalization data, we found that *PTBP1* contains a strongly supported colocalizing sQTL (H_4_PP: 0.99; # tissues: 9), associated with the exclusion or inclusion of exon 9 (**Figure 6B-C**). The sQTL junction mapped to 11 isoforms in the GENCODE annotation, but likely corresponds to the two major isoforms in hFOBs: *PTBP1-201* and *PTBP1-203*, which excludes and includes exon 9, respectively. Exon 9 resides in a linker region between RNA recognition motif (RRM) domains, and its skipping reduces the spacing between RRM domains and, consequently, reduces its ability to repress exons, including cryptic exons ^30,31^. Therefore, the mechanism of action for this sQTL may be the altered ratio of repressive (*PTBP1-203*) to non-repressive (*PTBP1-201*) isoforms, with higher ratios associated with decreased BMD (**Figure 6D**). This genetically-regulated isoform balance could, in turn, globally regulate the repression of *PTBP1* target exons during osteoblast differentiation. In support of this, we found that lead sQTLs for BMD revealed a clear enrichment in *PTBP1* binding sites (Fisher’s exact test, FDR = 0.019, in HepG2 cell), suggesting that both *PTBP1* trans-factor and its cis-regulatory targets are part of a splicing regulatory network that could mediate BMD.

### Long read proteogenomics to delineate sQTL-associated protein isoforms

The full-length transcriptome reference enables first-pass interpretation of sQTLs. But many sQTLs likely exert their effect on protein functions. Risk variants residing in GWAS loci likely modulate the particular ratios of protein isoforms, with a range of possible molecular functional effects, from overall loss of protein abundance from nonsense mediated decay (NMD) or degradation mechanisms, to gain of functions from production of multiple, functionally distinct protein isoforms from the same gene ^32,33^. However, to date, information on the protein isoforms of relevance has been largely unattainable. We recently developed a “long-read proteogenomics” pipeline ^15^, in which a full-length transcriptome is translated into ORFs to generate sample-specific protein isoform models. We employed this approach to generate high quality protein isoform models for osteoblasts. Onto these protein isoforms, we mapped colocalized sQTLs, finding mappings—defined as all cases in which a junction is wholly residing within a predicted coding sequence (i.e., ORF)—to 2,029 protein isoforms from 441 genes (**Supplemental notes 7 and 8**).

We systematically characterized the impact of sQTL-induced protein isoform changes (**Supplemental note 9**). Using a custom pipeline, Biosurfer (**Methods**), we compared groups of protein isoforms corresponding to non-risk (associated with increased BMD) versus risk (associated with decreased BMD) sQTLs, in effect, bioinformatically quantifying putative protein-level effects. Protein isoform sequences were compared using a custom hybrid alignment developed for the task, which aligns proteins based on their sequence, but with consideration of the underlying exon-intron structure and junctions corresponding to sQTLs. We performed pairwise comparisons between all risk and non-risk isoforms, weighted by their gene expression in hFOBs, and determined if, overall, the effect on the protein was one that led to a shift to transcripts predicted to undergo NMD (e.g., a possible lowering of protein abundance), a dramatic reduction in protein length, or if the protein models indicate a potential change in two functionally relevant proteins, as indicated by those proteins that are full-length with preserved domain structure.

We found that of the 809 sQTLs, only 74 likely represent a switch from a full-length protein product to a transcript undergoing NMD, by presence of a premature stop codon upstream of at least one junction ^34^ (**Table S3**). The remaining sQTL-protein-isoform groups involve a switch from one full-length protein to another, albeit with vast differences in length. We found 190 sQTLs that potentially lead to a truncated protein which may represent a sub functional form, or even a dominant negative ^35^ (**Table S4**). Although as previously observed ^36^ reduced length does not always correlate with reduced functional capacity ^36^. To our knowledge, this is the first proteome-scale dataset of protein isoforms associated with a clinical trait by mapping directly from an sQTL dataset.

### Distinct isoforms of *TPM2* have opposing effects on osteoblast differentiation and mineralization

Our analysis identified 441 genes and 2,029 protein isoforms potentially contributing to BMD. We next aimed to prioritize genes to select a set of candidate isoforms to test for a possible role in osteoblast differentiation and mineralization. We evaluated candidate genes primarily based on the strength of colocalization (H_4_PP) and the extent of sQTL sharing across tissues (reasoning that genes with sQTL observed in multiple tissues were more likely active in osteoblasts). We also evaluated additional data such as whether genes were the cause of a monogenic bone disease (**Tables S5 and S6**). We prioritized genes not previously implicated in the regulation of BMD.

Based on these criteria, we selected *TPM2* for further investigation. *TPM2* codes for beta-tropomyosin, which regulates the contractile machinery of muscle cells ^37^. TPM2 is a coiled-coil protein and a component of actin filaments, where it regulates the interaction of actin and myosin during muscle contraction ^38^. *TPM2* mutations have been linked to various muscle disorders, including nemaline myopathy, a congenital disorder characterized by muscle weakness and wasting ^44^. *TPM2* has not been experimentally demonstrated to play a role in osteoblast activity or BMD; however, we previously identified it as a strong candidate for the GWAS association it resides in using a network-based approach ^39^. Furthermore, in an independently mapped protein-protein interactome network (STRINGdb ^40^), TPM2 resides in a cluster enriched in bone relevant terms such as the Monarch ^41^ category “phenotype of bone density” (EFO:0003923, FDR = 1.68×10^−16^). In addition, quantitative serum proteomics of individuals with low and high BMD revealed TPM2 protein to be differentially expressed ^42^. Finally, in mice, *TPM2* is highly expressed in primary calvarial osteoblasts with an apparent lack of expression in other bone cells ^43^.

Based on our long-read RNA-seq data in hFOBs, four primary *TPM2* isoforms were expressed. They differ based on a pair of mutually exclusive exons (exons 6 and 7) and a pair of alternative last exons (exons 10 and 11), leading to the following four isoforms: TPM2-6-10 (*TPM2* isoform containing exons 6 and 10), TPM2-6-11, TPM2-7-10, and TPM2-7-11 (**Figure 7A**). We developed a targeted mass spectrometry method ^44^ for peptides specific to shared and alternative exons of *TPM2*, including exons 6, 7, 10, and 11 (**Supplemental note 10**), confirming the expression of TPM2 protein isoforms in hFOBs.

**Figure 7:**
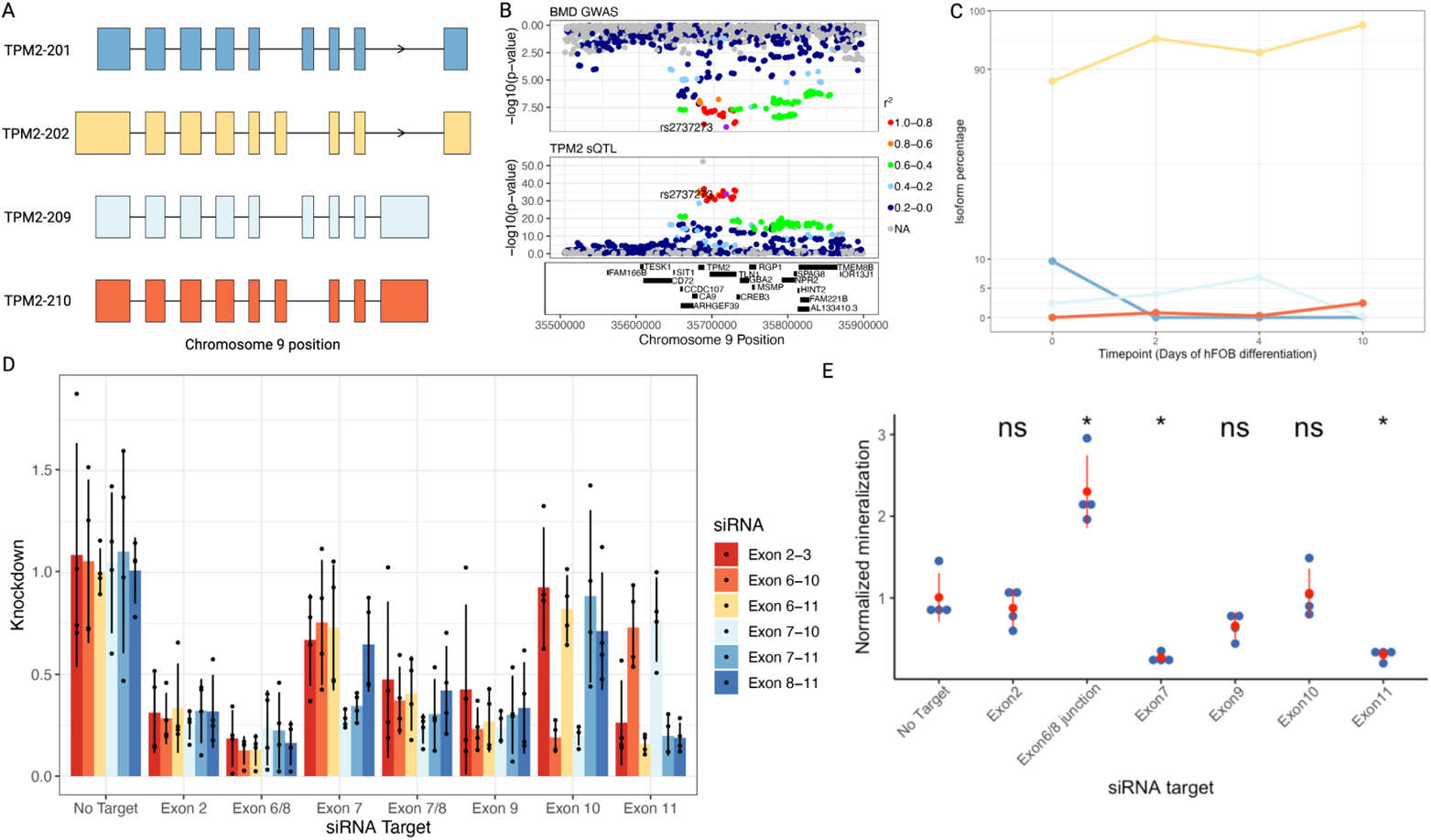
Different TPM2 isoforms show opposing effects on mineralization. A) Isoform models for *TPM2* isoforms expressed in hFOBs. B) Mirrorplot showing a significantly colocalizing sQTL around exons 6 and 7 in *TPM2* (H_4_PP = 0.98). C) Isoform percentages across hFOB differentiation timepoints highlighting *TPM2-202* as the major isoform. D) siRNA knockdown of isoforms containing target exon (x-axis). Each color represents an siRNA target (red = isoforms containing exon 2, orange = isoform containing exons 6 and 10, yellow = isoforms containing exons 6 and 11, sky blue = isoforms containing exons 7 and 10, light blue = isoforms containing exon 7 and 11, dark blue = isoforms containing exon 11). E) Quantification of nodule mineralization using alizarin red staining in hFOBs (*) represents significant T-Test comparison between no target control and the target exon while (ns) represents no significant difference.

For *TPM2*, we identified multiple colocalizing sQTLs (H_4_PP ≥ 0.75) that converge on the two sets of splicing events. The sQTLs for junctions surrounding exons 6 and 7 were observed in a wide-range of tissues (13 to 33 tissues); whereas those around exons 10 and 11 were observed strictly in brain and testis. (**Figure 7B**). The T allele of the lead GWAS SNP (rs2737273) in the locus is associated with a decrease in BMD and an increase in *TPM2* isoforms containing exon 6. Conversely, the C allele is associated with an increase in BMD and increase in isoforms containing exon 7. We also observed in brain and testis that the T allele was associated with increased exon 11 containing isoforms and a decrease in BMD; however, it is difficult to interpret the direction of effects of multiple sQTL due to the correlated nature of multiple splicing events and events occurring across tissues.

Based on the sQTL effect information, the restriction of *Tpm2* expression to osteoblasts in the mouse (using BioGPS)^43^, and its high expression in human osteoblasts (hFOBs) (**Figure 7C**), we hypothesized that exon 6 containing *TPM2* isoforms would be negatively associated with mineralization in hFOBs, with the opposite effect found for exon 7 containing isoforms. To test this hypothesis, we performed isoform-specific siRNA knockdown experiments in hFOBs followed by quantification of mineralized nodules (**Figure 7D and E**). First, we knocked down all isoforms of *TPM2* by targeting constitutively expressed exon 2 and exon 9 (**Figure 7D**), and for both experiments we observed no significant differences in mineralization relative to the control (T-Test, p = 0.51 and p = 0.09 respectively), demonstrating that knock down of all *TPM2* isoforms simultaneously does not result in a phenotype (**Figure 7E**). Next, we targeted *TPM2* isoforms containing exon 6 (targeting the junction between exons 6-8) (**Figure 7D**), and, strikingly, observed a significant increase in mineralization relative to the control (T-Test, p = 0.004) (**Figure 7E**). This siRNA moderately decreased the expression of all isoforms, though it did attenuate expression of exon 6 containing isoforms to the greatest degree (**Figure 7D**). On the other hand, knockdown of isoforms containing exon 7 (**Figure 7D**) resulted in a significant decrease in mineralization in accordance with our hypothesis (T-Test, p = 0.01) (**Figure 7E**). To comprehensively assess the effect of all *TPM2* splice events, we also targeted exons 10 and 11. We observed no change in mineralization upon knockdown of exon 10 containing isoforms (T-Test, p = 0.82). Surprisingly, knockdown of exon 11 containing isoforms (**Figure 7D**) decreased mineralization (T-Test, p = 0.015) (**Figure 7E**). These data suggest that disruptions in the ratios of the four *TPM2* isoforms have significant impact on mineralization in hFOBs. Most importantly, our results clearly demonstrate that different *TPM2* isoforms have distinct functions with respect to osteoblast activity and are likely regulators of BMD.

## Discussion

A wealth of GWAS and sQTL datasets has highlighted the widespread involvement of alternative splicing in patient disease risk ^3,5^, but how individual sQTLs affect downstream transcript and protein isoforms to mediate disease is largely unknown. Here, we present a broadly applicable method to increase the interpretability of such sQTLs. The method integrates sQTLs with long-read RNA-seq data to nominate full-length protein isoforms impacted by colocalized sQTLs, accelerating the path for functionally characterizing the molecular implications of sQTLs.

As a proof-of-principle, we applied our approach to BMD, the single strongest predictor of osteoporotic fracture ^45^. We identified thousands of candidate full-length isoforms potentially involved in the regulation of BMD, providing strong support for the hypothesis that splicing is a major mediator of genetic variation impacting bone, similar to other traits ^6,46,47^. However, our results extend these studies by providing the first resource to specifically connect transcript and protein isoform expression to biological processes and pathways impacting disease.

Our approach nominated isoforms predicted to influence BMD for the gene *TPM2. TPM2* splicing has not been linked to osteoblast differentiation or bone disease; however, point mutations in *TPM2* lead to muscle diseases such as the Escobar variant (characterized by skeletal defects including vertebral defects, bone fusion abnormalities and growth retardation) ^48^, nemaline myopathy ^49^, and atherosclerosis ^50^. Additionally, loss of *Tpm2.1* (the isoform containing both exons 6 and 11) in the mouse increases beta catenin levels ^51^, which is integral to osteoblast activity and bone formation ^52^. We demonstrated that isoforms containing exon 6 and exon 7 show opposing effects on osteoblast mineralization. Intriguingly, we also observed decreased mineralization upon knocking down isoforms containing exon 11. This was unexpected since the isoform with exons 6 and 11 was the most highly expressed and a relatively small decrease in exon 6 isoforms compared to all other isoforms significantly increased mineralization. These results suggest non-additive interactions among *TPM2* isoforms, and that osteoblast activity and mineralization must be highly dependent on isoform stoichiometries during differentiation.

Globally, among the hundreds of genes with colocalizing sQTLs, we identified an enrichment for genes involved in alternative splicing regulation. Both trans-acting splice factors and their cis-regulatory targets were affected, suggesting a possible convergence onto the same splicing network. For example, we found a colocalizing sQTL in *PTBP1*, a trans-acting splice factor that mediates alternative splicing by binding mainly to the polypyrimidine tract ^53^. Interestingly, *PTBP1* isoforms including exon 9 have been shown to bind upstream of and represses the inclusion of exon 7 in *TPM2*—another gene with colocalized sQTLs—leading to higher levels exon 6, the mutually exclusive exon partner of exon 7, in *TPM2* ^*28,31*^. Intriguingly, the genetic associations indicate that higher levels of *PTBP1* including exon 9 (lead SNP: rs2737273) and higher levels of *TPM2* including exon 6 (lead SNP: rs3215700) are associated with lower BMD. Together, this example of a *PTBP1*-*TPM2* splicing axis highlights the functional interconnectivity of splicing network components, which, in turn, can be genetically modulated by sQTLs.

Our work represents one “template” or version of integrating GWAS, sQTL, and long-read transcriptomics for characterizing genetically regulated protein isoforms. Few transcriptomic datasets are available for tissues directly relevant to BMD, such as bone cells, therefore, we obtained candidate splice sites by using GTEx sQTLs and we focused on sQTLs that are highly shared across tissues ^3^. To contextualize the GTEx sQTLs, we generated long-read RNA-seq in a model system directly relevant to bone to identify expressed isoforms. In this sense the long-read RNA-seq data provided a biological context “filter” to nominate the most relevant isoforms for functional validation.

Another variation of our approach is generating a comprehensive long-read transcriptome reference from a subset of the same samples used for short-read based sQTL discovery, which can serve as a higher accuracy isoform reference for aligning and assembling short-read RNA-seq reads. However, this approach is still subject to well-known limitations of short-read-based isoform characterization ^54^. Finally, as the throughput of long-read RNA-seq reaches that of traditional short-read methods, with recent developments from PacBio and ONT platforms, among others, it will become feasible to generate high quality long-read data at population scales that would enable direct discovery of isoform-level QTLs. It is also likely that variations of our approach could use diverse methods of detecting genetically-regulated splicing signals, beyond colocalization approaches, such as transcriptome-wide association studies (TWAS).

In this study, we identified hundreds of potentially causal isoforms in osteoporosis. However, limitations remain. One of the limitations was the use of non-bone sQTLs from GTEx ^3^, due to the lack of population scale transcriptomics datasets on bone or bone cells. Therefore, our analysis missed sQTL associations that are specific to bone. In addition, the use of sQTL from multiple non-bone tissues may have inflated the number of false positives due to colocalization signals that have no biological impact on bone. Though, we expect that the overall false positive rate was reduced based on the requirement that isoforms were expressed in human osteoblasts. Finally, we demonstrated the utility of our approach on a single cell type (osteoblasts), but future work should include long-read RNA-seq data from a more comprehensive collection of cell-types with relevance to bone (e.g., osteoclasts, and osteocytes).

Central to our method is the use of long-read-driven isoform models serving as a scaffold for interpreting colocalized sQTLs; therefore, the quality of transcriptome-wide maps remains critical for appropriate biological interpretation of isoform effects. Long-read RNA-seq technologies continue to evolve at a rapid pace, with concomitant need for evaluating their analytical metrics (e.g., accuracy, and quantitative precision). Recently, the Long Read Genome Annotation Consortium (LRGASP) community effort is aiming to comprehensively benchmark such metrics (Registered report at *Nature Methods*). At least some sources of biochemical, sequencing, and bioinformatic artifacts have been previously discussed ^55^. As it stands, no method yet exists for large-scale detection of isoforms at the protein level, therefore, for tractability, we employed a bioinformatic proteogenomics approach in which the transcript models are used as a proxy for inferring the putative protein isoform products, similarly to the reference annotation ^56^. Given the state of protein isoform information, all isoforms of analysis should be fully validated, as we have done with *TPM2* using MS-based proteomics. Even with full knowledge of protein isoforms expressed in disease relevant models, limitations arising from the nature of complex loci containing many distinct sQTLs and highly complex splicing, in terms of number of events and dependencies between distal splicing, may still be out of reach for straightforward interpretation without extensive functional validation.

In this study, we developed an integrative systems genetics approach to identify isoforms that are potentially responsible for the effects of BMD GWAS loci ^17^. This is the first study to our knowledge that directly incorporates long-read RNA-seq to systematically nominate protein isoforms that are potentially causal effectors of BMD. These results highlight the importance of genetically regulated alternative splicing to BMD and identifies hundreds of potential drug targets for osteoporosis and bone fragility. Our work should serve as a model for other researchers to increase the clinical utility of sQTL analysis across the spectrum of human diseases.

## Methods

### GWAS and sQTL analysis

eBMD GWAS summary statistics were downloaded from the GEnetic Factors for OSteoporosis Consortium (GEFOS) website using http://www.gefos.org/?q=content/data-release-2018 (accessed July 2021). The coordinates of the GWAS SNPs were updated from hg19 to hg38 using LiftOver within Bioconductor. GTEx V8 sQTL all association summary statistics data were downloaded from the GTEx Google Cloud Platform (GCP) using https://console.cloud.google.com/storage/browser/gtex-resources (accessed July 2021). The data were prepared as input for coloc ^20^ as follows: A list of genes with a start site that is within ∓200 Kb of each eBMD GWAS SNP was created. Using this list, a list of sQTL associations within ∓200 Kb of each of these genes was created for each GTEx tissue. coloc.abf was used with this input (all the GWAS SNPs and all the sQTL data within ∓200 Kb of each gene start site). In order for a junction to be considered significant (junction with a colocalizing sQTL), it must have coloc ^20^ H_4_PP of ≥ 0.75.

### Osteoblast differentiation

hFOB 1.19 (American type culture center [ATCC], Manassas, VA; CRL-11372) were grown and differentiated with RNA isolated on days 0, 2, 4 and 10 exactly as outlined in Abood et al. ^57^. Briefly, hFOB 1.19 cells (American Type Culture Collection [ATCC], Manassas, VA, USA; #CRL-11372) were cultured at 34°C and differentiated at 39.5°C as recommended with the following modifications. Growth media: minimal essential media (MEM; Gibco, Grand Island, NY, USA; 10370-021) supplemented with 10% fetal bovine serum (FBS; Atlantic Biologicals, Morrisville, NC, USA; S12450), 1% Glutamax (Gibco; 35050-061), 1% Pen Strep (Gibco; 15140-122). Differentiation media: MEM alpha (Gibco; 12571-063) supplemented with 10% FBS, 1% Glutamax, 1% Pen Strep, 50 μg/μL Ascorbic Acid (Sigma-Aldrich, St. Louis, MO, USA; A4544-25G), 10mM beta-Glycerophosphate (Sigma-Aldrich; G9422-100G), 10 nM Dexamethasone (Sigma-Aldrich; D4902-25MG). RNA was isolated from ∼0.5 × 10^6^ cells at days 0, 2, 4, and 10 of differentiation as recommended (RNAeasy Minikit; QIAGEN, Valencia, CA, USA; 74106). Mineralized nodule formation was measured by staining cultures with Alizarin Red (40 mM, pH 5.6; Sigma-Aldrich; A5533-25G). Reported results were obtained from three biological replicate experiments for days 2, 4, and 10, and two biological replicates for day 0 of differentiation.

### Long-read RNA-seq data collection and sequencing

RNA was extracted from three biological replicates of hFOB cells at days (0, 2, 4, and 10) using Qiagen RNeasy mini kit. Extracted RNA was used to create full length cDNA using the NEBNext® Single Cell/Low Input cDNA Synthesis & Amplification kit (New England Biolabs Ipswich MA Lot#10078130). Following the PacBio Iso-seq protocol, first and second strand cDNA were synthesized using NEB dT oligo and the PacBio Template Switching Oligo. For the cDNA amplification 14 cycles of PCR were performed followed by a Pronex bead cleanup (Promega Corporation Madison WI, Lot #NG103A). The amplified and purified cDNA were QCed using the Bioanalyzer DNA 12000 kit. The samples were then sent to the Smith lab at the University of Louisville for long-read PacBio sequencing.

Approximately 300 ng of cDNA was converted into a SMRTbell library using the Iso-Seq Express Kit SMRT Bell Express Template prep kit 2.0 (Pacific Biosciences). This protocol employs bead-based size selection to remove low mass cDNA, specifically using an 86:100 bead-to-sample ratio (Pronex Beads, Promega). Library preparations were performed in technical duplicate. We sequenced libraries using 11 SMRT cells on the Sequel II system using polymerase v2.1 with a loading concentration of 85pM. A 2-hour extension and 30-hour movie collection time was used for data collection. The “ccs” command from the PacBio SMRTLink suite (SMRTLink version 9) was used to convert raw reads (∼22 million) into circular consensus sequence (CCS) reads. CCS reads with a minimum of three full passes and a 99% minimum predicted accuracy (QV20) were kept for further analysis.

### Long-read RNA-seq differential analysis

All differential statistical analyses in long-read data were performed using tappAS ^23^. The input files for tappAS are the raw expression matrix obtained from cDNA Cupcake, the full-length transcriptome reference file generated from IsoAnnot, and a design matrix for time-series analysis. Within tappAS, maSigPro ^58^ was chosen for differential transcript expression using the following parameters: polynomial degree = 3, alpha = 0.05, R^2^ cutoff = 0.7, and max K clusters = 10 however “mclust” ^59^ was used to ensure an optimal number of clusters. Differential isoform usage analysis was performed within tappAS using maSigPro with the following parameters: polynomial degree = 3, alpha = 0.05.

### Quantitative mass spectrometry-based proteomics

#### Protein extraction, quantitation, and digestion

hFOB cells used for proteomic mass spec analysis were isolated by removing media from a 10 cm plate of cells (∼5 × 10^6^) and subsequently washing with 5 ml of PBS. The cells at day 0, were treated with 2 ml 0.05% trypsin/EDTA for 5 minutes at 34°C, triturated, pelleted at 1000xg for 5 minutes, washed with PBS, pelleted, resuspended in 1.5 ml PBS, transferred to a 1.5 ml microfuge tube, pelleted with the pellet snap frozen and stored at -80°C. The cells at day 2 and 4 were treated with 2.5 ml, 8 mg collagenase (Gibco 17018-029) / ml HBSS (Hanks Balanced Salt Solution [HBSS], Gibco 14025-092, supplemented with 4mM CaCl) for 15 minutes at 37°C followed by the addition of 2.5 ml 0.25% trypsin/EDTA with incubation at 37°C continued for an additional 15 minutes. Cells were subsequently treated as outlined for day 0 cells. The cells at day 10 were initially incubated with 5 ml 60 mM EDTA, pH 7.4 (prepared in PBS) for 15 minutes at RT. The EDTA solution was removed and cells washed with HBSS. The cells were then subsequently treated as outlined for day 2 and day 4 cells

Harvested hFOB cells, approximately 8-10 million cell count each, were pelleted in triplicate for four different time points: day 0, day 2, day 4, day 10. The twelve samples were frozen at -80°C until lysis. Each pellet was lysed according to the Filter Aided Sample Preparation (FASP) protocol adapted from ^60^. Lysis buffer was changed to 6% SDS, 150 mM DTT, 75 mM Tris-HCl. To each pellet, an aliquot lysis buffer equal to 2x the pellet volume was added and probe-sonicated to lyse the cells and shear the nucleotide material. Sonication continued for 1-5 minutes until the sample was clear and no longer viscous. The lysates were then incubated at 95°C for 2.5 minutes. Protein quantitation was estimated by BCA assay to be approximately 500-4000 ug per lysate. Aliquots equivalent to 80 ug per sample were used for FASP and buffer exchanged into 200 mM EPPS pH 8.5. A technical replicate of day 10 replicate C was prepared as well as an unrelated Jurkat lysate sample, resulting in a total of 14 samples for proteolytic digestion. Digestion was performed as per Navarrete-Perea et al. ^61^ with Lys-C overnight, followed by trypsin for six hours, using a 1:100 enzyme-to-protein ratio.

#### Tandem Mass Tag (TMTpro) labeling

Reagents from the TMTpro 16plex isobaric label reagent set A44522 (ThermoFisher, Waltham, MA) were used for labeling each of the 14 samples (TMTpro-133C and TMTpro-134N were not needed). Per protocol, 20 uL of anhydrous acetonitrile was added to each tube of 0.5 mg dry TMTpro reagent and allowed to incubate at room temperature for 5 minutes with occasional vortexing. The entirety of each TMTpro vial was added to its corresponding digest sample and allowed to incubate at room temperature for one hour, vortexing every 10 minutes, followed by a final centrifugation of the tubes.

#### C18 desalting, label efficiency check

Aliquots of 2 uL (1.5%) of each digest were combined and desalted using EasyPep Mini C18 desalting resin (ThermoFisher, Waltham, MA). The remaining sample was kept frozen at -80℃until the next day. Eluted sample was dried via speed vac and reconstituted in 6 uL of 0.1% formic acid. The entire sample was injected for LC-MS/MS analysis to check labeling efficiency and mixing ratios among the 14 labeled samples. After check analysis, the remaining sample from all digests were brought back to room temperature, quenched according to TMTpro protocol, and mixed according to normalized ratios determined from this check analysis (see below).

#### LC-MS/MS check analysis

Desalted sample was analyzed by nanoLC-MS/MS using a Dionex Ultimate 3000 (Thermo Fisher Scientific, Bremen, Germany) coupled to an Orbitrap Eclipse Tribrid mass spectrometer (Thermo Fisher Scientific, Bremen, Germany). Six microliters (estimated 1ug) were loaded onto an Acclaim PepMap 100 trap column (300 um x 5 mm x 5 um C18) and gradient-eluted from an Acclaim PepMap 100 analytical column (75 um x 25 cm, 3 um C18) equilibrated in 96% solvent A (0.1% formic acid in water) and 4% solvent B (80% acetonitrile in 0.1% formic acid). The peptides were eluted at 300 nL/min using the following gradient: 4% B from 0-5 minutes, 4-10% B from 5-10 minutes, 10-35% B from 10-60 minutes, 35-55% B from 60-70 minutes, 55-90% B from 70-71 minutes, and 90% from 71-73 minutes.

The Orbitrap Eclipse was operated in positive ion mode with 2.1kV at the spray source, RF lens at 30% and data dependent MS/MS acquisition with XCalibur version 4.3.73.11. MS data acquisition was set up according to the existing method template, “TMT SPS-MS3 RTS”. Positive ion Full MS scans were acquired in the Orbitrap from 400-1600 m/z with 120,000 resolution. Data dependent selection of precursor ions was performed in Cycle Time mode, with 2.5 seconds in between Master Scans, using an intensity threshold of 5 × 10^3^ on counts and applying dynamic exclusion (n=1 scans for an exclusion duration of 60 seconds and ∓10 ppm mass tolerance). Monoisotopic peak determination was applied and charge states 2-8 were included for CID scans (quadrupole isolation mode; rapid scan rate, 0.7 m/z isolation window, 32% collision energy, AGC standard). MS3 quantification scans were performed when triggered by the real-time search (RTS) algorithm. MS3 (HCD) scans were collected in the Orbitrap with 50,000 resolution, 50% collision energy, AGC target of 300%, and automatic maximum inject time mode for a maximum of 10 SPS precursors per cycle.

#### Sample preparation for targeting *TPM2* peptides

Replicates “A” and “C” of each of the following TMTpro-labeled samples were mixed according to normalized ratios determined from the check analysis: Days 0, 2, 4, 10. In addition, one technical replicate for Day 10 was included, as well as one Jurkat sample (used as a negative control). An estimated 6.6 ug from each sample was mixed together, for a total of 66 ug. The pooled sample was subjected to C18 desalting using EasyPep Mini C18 desalting resin (ThermoFisher, Waltham, MA), reconstituted with 66 uL 0.1% formic acid for 1 ug/uL concentration, and used for tMS2 targeting analysis without further fractionation.

#### LC-MS/MS: tMS2 analysis

The resulting peptides were analyzed by nanoLC-MS/MS using a Dionex Ultimate 3000 (Thermo Fisher Scientific, Bremen, Germany) coupled to an Orbitrap Eclipse Tribrid mass spectrometer (Thermo Fisher Scientific, Bremen, Germany). One microliter (estimated 1 ug) was loaded onto an Acclaim PepMap 100 trap column (300 um x 5 mm x 5 um C18) and gradient-eluted from an Acclaim PepMap 100 analytical column (75 um x 25 cm, 3 um C18) equilibrated in 96% solvent A (0.1% formic acid in water) and 4% solvent B (80% acetonitrile in 0.1% formic acid). The peptides were eluted at 300 nL/minute using the following gradient: 4% B from 0-5 minutes, 4-28% B from 5-210 minutes, 28-50% B from 210-240 minutes, 50-95% B from 240-245 minutes and 95% B from 245-250 minutes.

#### tMS2 mode

A total of 22 peptides from source protein *TPM2* were selected for tMS2 targeting. Charge states z = 2 and z = 3 were selected for each peptide for a total of 44 target m/z values (**Table S7**). The Orbitrap Eclipse was operated in positive ion mode with 2.1kV at the spray source and RF lens at 30% with XCalibur version 4.5. In tMS2 mode, no MS1 scans were acquired and MS2 scans were isolated in the quadrupole with a 1.6 m/z isolation window and fragmented using HCD with 30% collision energy. Standard AGC was used and fragment ions were detected in the Orbitrap with 30,000 resolution. Retention time scheduling was not used.

#### Mass spectrometric data analysis Quantification

Impurity correction factors were included for reporter ion intensities, based on the quality control information provided with the TMTpro reagent kit. Reporter ion abundances above a minimum S/N of 2 and below a co-isolation threshold of 70% were summed across peptide spectral matches (PSM) to calculate peptide abundance. Normalization mode was set to “Total Peptide Amount” and scaling mode was “On All Average” so that the average abundance per protein and peptide was normalized to 100. Quantification rollup parameters were set to “Protein Abundance Based” protein ratio calculation, allowing for a maximum 100-fold change in abundance, with low abundance resampling imputation and ANOVA (individual proteins) hypothesis testing.

#### Manual inspection of targeted TPM2 peptides

Skyline viewer program ^62^ was used to view and validate MS2 spectra for target peptides. For quantitative analysis, the ion counts of individual reporters ions were recorded for each target peptide MS2 spectrum using FreeStyle raw file viewer.

### Long-read RNA-seq analysis pipeline

Long-read RNA-seq processing was performed using the Isoseq3 workflow (https://github.com/PacificBiosciences/IsoSeq). Primer removal and demultiplexing of cells was performed using “lima”. Next, the demultiplexed samples were refined by keeping only transcripts with poly-A-tails and removing any concatemers using the module “refine”. Following that the full-length non-chimeric poly-A containing reads were clustered into isoforms using the module “cluster” and then aligned using the PacBio read compatible minimap2 ^63^ aligner. Finally the clusters of isoforms were collapsed into non-redundant isoforms using the “collapse” module. A raw matrix of expression was generated using cDNA Cupcake’s “demux_isoseq_with_genome.py” module. Post Isoseq3, SQANTI3 ^64^ was used to classify isoforms into five categories: FSM (Full splice match), ISM (Incomplete splice match), NIC (Novel in catalog), NNC (Novel not in catalog), and genic. Following that, IsoAnnot was used to generate a full-length transcriptome reference from long-read data.

### Long-read proteogenomics analysis pipeline

We relied heavily on the published long-read proteogenomics pipeline ^15^. In short, open-reading frame identification steps used CPAT ^65^ to assess coding potential of isoforms, followed by generation of predicted ORFs using the “ORF_calling” module. Next, we generated the CDS GTF file to obtain the list of proteoforms using “make_pacbio_CDS” and “refinement” modules. We performed NMD and truncation analyses using Biosurfer (https://github.com/sheynkman-lab/Biosurfer_BMD_analysis). Briefly, each ORF containing an sQTL junction is compared against all ORFs not containing this sQTL and both length differences and the NMD rule are applied (See **supplemental note 9**).

### Experimental validation of *TPM2* in hFOBs

#### siRNA Knockdown; general information

hFOB cells used in the siRNA knockdown experiments were transfected within five days of thawing from liquid nitrogen storage. Briefly, cells were seeded at 75,000 cell/well of a 24 well plate and transfected within 24 hours after plating. In a typical experiment, 10 wells were seeded for each siRNA treatment (7 and a No Target control, see Table S8) plus a ‘not transfected’ control as well as 4 wells for RNA collected on the day of transfection (94 wells in toto). Four wells of each treatment were used for determining the amount of mineral formed at differentiation Day 10 with the remaining six wells used for RNA which was isolated at various times during the course of the experiment.

#### siRNA transfection procedure and conditions

Within 24 hours after plating, cells were transfected with siRNAs using Lipofectamine LTX reagent (Invitrogen ref# 15338100) following the manufacturer’s recommended procedure, with minor modifications. Briefly, between 16-18 hours after plating, 2.5 ul Lipofectamine LTX was mixed with 37.5 ul pre-warmed Opti-MEM (Gibco, ref # 31985-070) per well of cells transfected. In parallel, 1 ul of 5 uM siRNA was mixed with 37.5 ul pre-warmed Opti-MEM per well. After a minimum of 5 minutes incubation time at room temperature (RT), the Lipofectamine mix and siRNA mix were combined, mixed and allowed to incubate for a minimum of 15 minutes at RT. After this time period, the ∼75 ul Lipofectamine/siRNA mixture was diluted into 500ul pre-warmed Opti-MEM/well, mixed, and applied to a well of cells after growth media was removed. Cells were placed back in the incubator (temperature=34°C) for 5-7 hours at which time the Opti-MEM transfection mix was removed and replaced with 500 ul/well of pre-warmed growth media after the cells were washed with pre-warmed DPBS (Dulbeco’s Phosphate Buffered Saline, Gibco, ref# 14190-144).

#### Osteoblast Differentiation

Twenty-four hours after transfection (48 hours after plating) growth media was removed, cells washed with 0.5 ml/well pre-warmed DPBS, 0.5ml/well pre-warmed differentiation media added and cells placed at 39.5°C. Differentiation media was replaced every other day after cells were washed with DPBS. On the tenth day, cells were washed 3x with 1 ml DPBS/well, fixed with 0.5 ml 10% Buffered Formalin Phosphate (Fisher #SF100-4) for 15 minutes at RT, washed three times with 1 ml/well water and stained with 400 ul/well 40 mM Alizarin Red, pH 5.6 (with NH4OH; Sigma A5533-25G) for 25 minutes at RT after which the stain was removed and cells washed 10x for 10 minutes each with 1 ml/well water. After images were scanned, the amount of alizarin red bound to the formed mineral was quantified by eluting in 2 ml/well 5% (v/v) Perchloric Acid (HClO4, Sigma-Aldrich 311413-500ML) and incubated for 20 minutes with shaking at RT. The amount of alizarin red bound for each sample/treatment was determined from the absorbance at 405 nm wavelength of the eluent along with standards prepared from the alizarin red staining solution appropriately diluted into 5% HClO4 and was expressed in units of nmol alizarin red bound.

#### RNA isolation and cDNA preparation

RNA was isolated using the Qiagen RNase minikit (cat# 74106) following the manufacturer’s protocol. Briefly, media was removed from cells and washed 3x 1 ml/well DPBS and subsequently lysed in 400 ul/well RLT buffer containing 40 mM dithiothreitol (DTT) with mild shaking for 10 minutes, transferred to a microfuge tube and stored at -80°C until processing. At the time of processing, RNA was isolated from thawed lysates exactly as outlined. RNA was immediately DNased after isolation Applied Biosystem’s TURBO DNA-free kit (Invitrogen ref# AM1907) following the manufacturer’s protocol. DNased samples were stored -80°C. cDNA was prepared from thawed samples by initially determining the RNA concentration with a Qubit 4 fluorometer and the RNA HS assay kit (Thermo Fisher ref# Q33226 and Q32855, respectively) following the manufacturer’s protocol. Random primed cDNA was synthesized from 1 ug DNased RNA using Applied Biosystems High Capacity Reverse Transcription kit (cat# 4368813).

#### Determining the extent and duration of siRNA knock down with real time PCR

The relative abundance of different transcriptional isoforms of the Tropomyosin 2 gene (*TPM2*) was determined in duplicates for each sample/treatment from four separate experiments two days after siRNA transfection. Each reaction contained ∼10 ng cDNA, 800 nM each primer (see **Table S9**), 0.5X Power Up SYBR Green Master mix (Applied Biosystems ref# 100029285) in a 10ul reaction and amplified in a QuantStudio 5 Real-Time PCR system (Applied Biosystems, A28135) under these cycle conditions (50℃(2 minutes), 95℃(2 minutes); ([95℃(1 second), 60℃(30 seconds)] for 40 cycles); melt curve (95℃(1 second), 60℃(20 seconds), 95℃(1 second)). Relative quantification of the different transcriptional isoforms was determined by the 2 exp (–delta delta C(T)) method ^66^ using the geometric mean of the C(T) values of *CCDC47* and *CHMP2A* as the reference genes. Briefly, the cycle number in which half of the final amount of product produced (Cq/C(T)) is determined following the manufacturer’s recommendations. The C(T) for each sample/reaction for each *TPM2* primer pair is subtracted from the geometric mean of the C(T) of primer pairs for the genes *CCDC47* and *CHMP2A* for the same sample (ΔC(T)). Finally, the delta C(T) for a given primer pair for the ‘No Target Control’ sample is subtracted from the siRNA knockdown samples C(T)s for that particular primer pair resulting in the ΔΔC(T) (ddCT). The reported values are 2 ^(–)ddC(T)^.

## Acknowledgments

Research reported in this publication was supported in part by the National Institute of Arthritis and Musculoskeletal and Skin Diseases of the National Institutes of Health under Award Numbers R01AR071657 and R01AR077992 to CRF. AA was supported in part by a National Institutes of Health (NIH) Biomedical Data Sciences Training Grant T32LM012416. National Library of Medicine R01LM014017. We thank the IMPC for accessibility to BMD data in knockout mice (www.mousephenotype.org). We thank the donors in the Genotype-Tissue Expression (GTEx) Project and their families. GTEx was supported by the Common Fund of the Office of the Director of the National Institutes of Health, and by NCI, NHGRI, NHLBI, NIDA, NIMH, and NINDS. We also acknowledge Leon Sheynkman and Ben Jordan for their advice and technical knowledge.

## Author Contributions

Abdullah Abood: Conceptualization; formal analysis; investigation; methodology; validation; writing – original draft; writing – review and editing. Larry Mesner: Data curation; formal analysis; investigation; methodology. Erin Jeffery: Data curation; formal analysis; investigation; methodology. Mayank Murali: Formal analysis.

Micah Lehe: formal analysis. Jamie Saquing: Consultation. Charles R Farber: Conceptualization, Funding acquisition; writing – review and editing. Gloria Sheynkman: Conceptualization, Funding acquisition; writing – review and editing.

## Data Availability Statement

The raw data that support the findings of this study are openly available in Gene Expression Omnibus (GEO) [GSE224588]. All analysis code is available on GitHub https://github.com/aa9gj/Bone_proteogenomics_manuscript. Processed and input data is stored in Zenodo https://doi.org/10.5281/zenodo.7603851. Additionally, we have created a UCSC browser for dissemination of results https://tinyurl.com/Proteogenomics-browser.

---

**Figure S1:**
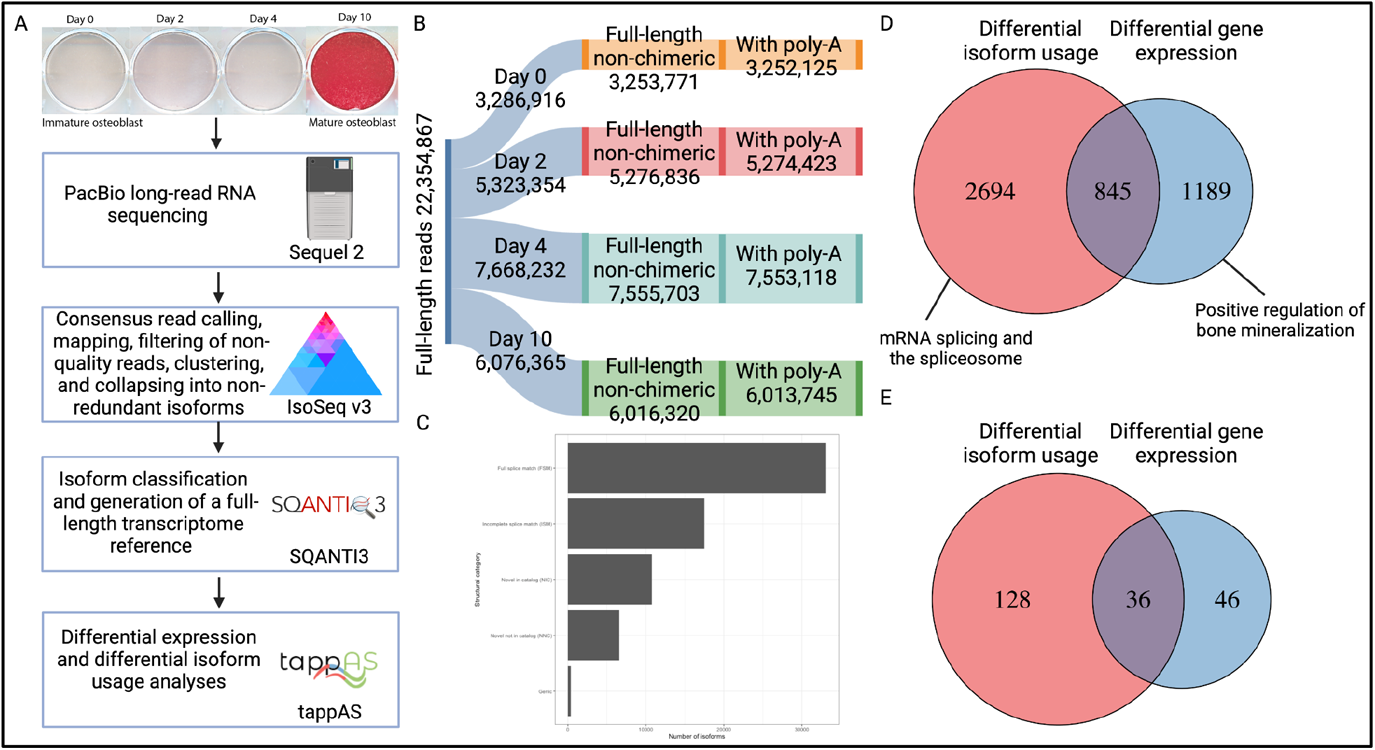
Long-read RNA-sequencing. A) Analysis pipeline for Long-read RNA-seq performed in hFOBs across 4 differentiation timepoints (Day 0, 2, 4, 10). Red color indicates mineralized nodules stained with alizarin red. B) Distribution of long-read RNA-seq across differentiation timepoints. C) Isoform classification. D) Venn diagram of the number of genes showing differential isoform usage and genes that are differentially expressed across hFOB differentiation along with representative Gene Ontology (GO) terms. E) Venn diagram showing genes with colocalizing sQTLs showing differential isoform usage and differential expression across hFOB differentiation.

**Figure S2:**
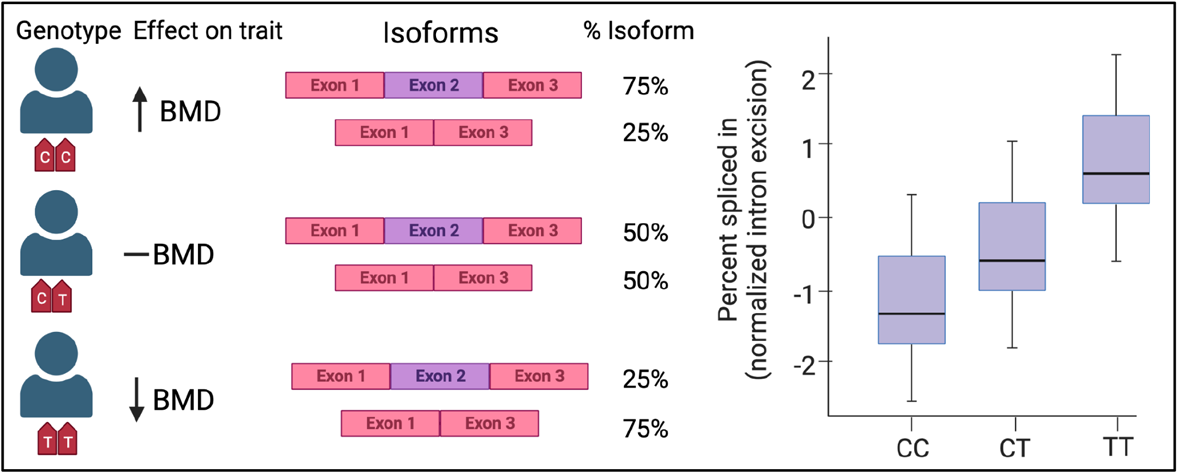
Mock illustration to determine how each isoform impacted BMD. The TT genotype is associated with a decrease in BMD (a result obtained from GWAS summary statistics). The same genotype is also associated with increased normalized intron excision ratio (from sQTL summary statistics). An increase in normalized intron excision ratio can be interpreted as an increase in the presence of the exon-exon junction. Therefore, we can hypothesize that genotype TT is associated with isoforms not containing exon 2 in this example which in turn is associated with a decrease in BMD. On the other hand, we can conclude that isoforms with exon 2 are associated with an increase in BMD.

