## Supplemental text for "Long-read proteogenomics to connect disease-associated sQTLs to the protein isoform effectors of disease"

### Supplemental note 1: Bayesian colocalization identified potential causal junctions in BMD

We performed Bayesian colocalization analysis with coloc [1] using the largest BMD GWAS [2] and existing sQTLs across 49 tissues from the GTEx project [3]. Overall, we found 820 genes with colocalizing sQTLs ($H_4PP \geq 0.75$) (total number of associations = 6,889) (**Table S10 and S11**). The number of unique junctions which we define as junctions with a unique donor-acceptor site pair regardless of the tissue where colocalization is reported is 2,043. We focused our study on protein-coding genes (GENCODE v38, gene "protein-coding") and their isoforms. The majority of sQTLs (6,391 of 6,889; ~93%) affect protein-coding genes (732 of 820; 89%) (**Table S12 and S13**), a finding similar to other studies [4]. The number of unique junctions is 1,863 found in 732 protein-coding genes (89% of total genes). Over half the junctions (51%) with colocalizing sQTL showed an $H_4PP$ of 0.90 or above.

### Supplemental note 2: Characterization of the full-length transcriptome in hFOBs using long-read RNAseq

We generated deep coverage long-read RNA-seq data across osteoblast differentiation using hFOB cell line. Long reads were collected in biological duplicate on day 0 and biological triplicate for days 2, 4, and 10, with a total of 22 million full-length reads obtained. A gradual increase in mineralization confirmed differentiation into functionally mature osteoblasts (see **Methods**, **Figure S1A-B**). Isoforms are considered detected if expressed in all replicates of at least one time-point and comprising a minimum of 1% isoform fractional abundance for the gene. We detected 68,326 transcript isoforms from 12,068 genes. The number of isoforms with a full-splice match (FSM) when compared to the GENCODE database is 33,106 (48% of total) while those with an incomplete splice match (ISM) is 17,482 (~26% of total). The number of novel isoforms is 17,738 (~26%) (**Figure S1C**). Of the novel isoforms, 10,793 (61%) arose from new combinations of known splice sites (NIC; novel in-catalog), whereas 6,580 (39%) arose from at least one novel splice donor or acceptor (NNC; novel not in-catalog) (**Figure S1C**). The median length of novel isoforms is 2,123 nt (mean = 2,268 nt). The median length of known isoforms is 2,055 nt (mean = 2,214 nt).

### Supplemental note 3: Mapping sQTLs colocalized with BMD onto the osteoblast-specific full-length transcript reference

The GTEx sQTLs are identified from non-bone tissues. Therefore, we sought to connect BMD-associated sQTLs directly to their source isoforms for proper interpretation of molecular effects. Accordingly, we proceeded to map the colocalized sQTLs onto the transcriptome of differentiating osteoblasts. The number of unique junctions with colocalizing sQTLs that are also observed in hFOBs is 836 (45% of colocalized junction in protein-coding genes); these junctions exactly map (donor-acceptor coordinates) to 459 protein-coding genes which in total have 2,349 isoforms (700 novel; ~30%). These genes are found within 362 associations (of the 1,103 BMD GWAS associations, 33%), with 221 lead associations harboring one sGene and 141 harboring more than one sGene (**Table S2**). Majority of the 836 junctions are found in known isoforms, specifically 383 (46%) are found in known isoforms only, and 350 (42%) are found in both known and novel isoforms. Strikingly, 103 (12%) of sQTLs mapped to novel isoforms. In order to confirm that the novel sQTL are in fact novel and are not a product of isoforms not expressed in hFOBs, we also mapped them to GENCODE v38. There were 73 sQTL (of 103; ~71%) that can be explained by known isoforms within GENCODE v38.

### Supplemental note 4: Identifying full-length isoform from colocalized sQTLs implicated in osteoblast differentiation

We next asked whether we see an enrichment of the genes with colocalizing sQTL for those undergoing splicing regulation in hFOBs differentiation. Over one third of the genes with colocalizing sQTLs show differential isoform usage (164/459; ~36% of genes with a colocalizing sQTL). Genes that show DIU are enriched in colocalizing sQTLs (Fisher's exact test, $p = 0.003$), indicating the relevance of the disease model (osteoblasts) and studying splicing in this process. On the other hand, we report 82 genes with colocalizing sQTL were differentially expressed (82/459; ~18%). However, genes that show DE are not enriched in colocalizing sQTLs (Fisher's exact test, $p = 0.57$). The number of genes with colocalizing sQTL that show both DE and DIU is 36 genes (**Figure S1E**).

### Supplemental note 5: Identification of high priority potential causal variants relevant to BMD

We aimed to provide a potential mechanism by which variants mediate splicing of introns. One way to investigate this is by characterizing the lead variants within junctions with colocalizing sQTLs. The total number of unique lead variants (lowest p value in the locus) associated with colocalized junction in hFOBs is 1,573 potentially regulating the splicing of these genes (as some lead SNPs have varying p-values due to tissue differences).

Among the colocalized sQTLs, we found that for over half of them (875/1,573, or 56%), the lead SNP or their proxy SNP in high linkage disequilibrium ($LD \geq 0.8$) resides directly within the associated intron **(Table S14)**, suggesting a potential regulatory mechanism associated likely to splice machinery. These proportions are similar to those reported for other complex diseases [5].

Next, we investigated whether these lead SNPs (or their proxy) are found within canonical splice sites (5' splice-donors and 3' splice-acceptors) in known and novel isoforms within hFOBs. We investigated whether these sQTLs are impacting unannotated splice sites, and two novel isoforms belonging to *DHRS12* (rs2296028: exon 8 acceptor site) and *PGS1* (rs11656568; exon 2 donor site) respectively showing disruptions to these sites (**Table S15**). This low number of SNPs that lie within these sites is expected as their disruption may be strongly deleterious [6].

### Supplemental note 6: Splice factors with colocalizing sQTLs

We obtained splice factor binding sites from CLIPdb in ENCODE [7]. We performed enrichment analysis of the lead sQTLs within the splice factor binding sites. We see significant enrichment of lead sQTLs within 34 of the 44 splice factors (**Table S16**). Of these, eight show differential expression across hFOB differentiation time points and 13 show differential isoform usage.

The splice factor hnRNPM shows differential expression across differentiation time points and a colocalizing sQTL. This gene has not been implicated previously in the regulation of BMD in human or mouse studies. However, evidence suggests that hnRNPM is implicated in an alternative splicing program Ewing sarcoma cells, which are aggressive tumors of bone and soft tissues [8]. Our results show three junctions with colocalizing sQTLs leading to the production of long and short isoforms of the gene.

### Supplemental note 7: Resource of candidate protein isoforms that mediate BMD

We compiled all the information on sQTL-linked protein isoforms, including their predicted risk status, creating the Proteoform for BMD Resource (PBR) (**Table S17**). We supplemented each isoform with additional evidence that may be pertinent to follow up studies. In terms of the relevance of the isoform in BMD and bone traits, we considered two major categories of evidence: i) Literature evidence pertaining to a novel role of the gene in bone processes and ii) data-driven evidence to contextualize our results. For the first category, we investigated whether the genes reported were implicated previously in bone monogenic disease (26 genes of 1,088; ~3%), have been previously identified as genes with colocalizing eQTLs in Al-Barghouthi et al. [9] (147 genes of 512; 32%), have been shown to influence bone strength in Diversity Outbred mice [10] (32 genes of 1,370; ~3%) or have been shown to disrupt BMD in IMPC [11] (14 genes of 371; ~4%). For our data-driven evidence, we leveraged differential isoform usage across hFOB differentiation (164 genes), extent of sQTL sharing across GTEx tissues, the strength of $H_4$PP, and the strength of the effect size (results are reported in **Tables S5 and S6**).

### Supplemental note 8: Mock example to illustrate connecting colocalized sQTLs to isoforms

We illustrate a toy example in **Figure S2**, To determine how each isoform impacted BMD, we cross referenced the effect size of the lead GWAS SNP with the directionality and magnitude of the slope in the same SNP from the GTEx sQTL data. In this example, the TT genotype is associated with a decrease in BMD (a result obtained from GWAS summary statistics). The same genotype is also associated with increased normalized intron excision ratio (from sQTL summary statistics). An Increase in normalized intron excision ratio can be interpreted as an increase in the presence of the exon-exon junction. Therefore, we can hypothesize that genotype TT is associated with isoforms not containing exon 2 in this example which in turn is associated with a decrease in BMD. On the other hand, we can conclude that isoforms with exon 2 are associated with an increase in BMD.

### Supplemental note 9: Nonsense mediated decay (NMD) and protein truncation prediction

To carry out the NMD and protein truncation analysis, we considered each colocalized sQTL and all the transcript isoforms of a gene to which the junction associated with the sQTL maps. All possible combinations of transcript isoforms from the sQTL are generated by pairing one isoform from the 'lacking' set with another from the 'containing' set. In this way, each resulting pair comprises one transcript isoform that lacks a particular exon and another that contains it. We then assigned weights to each pair of transcript isoforms, based on their average abundance values across the four time points. The weights of each pair were computed by multiplying the average abundance of the containing isoform with the average abundance of the lacking isoform and dividing by the product of the total abundance of all transcript isoforms in the 'containing' set and the total abundance of all transcript isoforms in the 'lacking' set. For example, *ST7L* gene transcript *PB.1181.14* (abundance = 10) in the 'containing' set and *PB.1181.36* (abundance = 50), *PB.1181.49* (abundance = 20) in the 'lacking' sets respectively. Then the weight for *PB.1181.14-PB.1181.36* isoform pair would be 500/700 or 0.71.

We investigated whether the colocalized sQTLs affect the isoform pairs involving one isoform predicted to undergo nonsense-mediated decay. For each 'lacking'-'containing' isoform pair, If the ORF has a stop codon located at least 50 base pairs upstream of the last splice site in the mature transcript (i.e. at the beginning of the last exon), it is considered a candidate for nonsense-mediated decay (represented as a boolean value, True if yes, False if not). Based on this identification, we assign an 'NMD status' as a product of the weight of that isoform pair and boolean value (1 if True, 0 if False) returned from predicting the NMD. Resulting in 'NMD status' either as 0 which would mean that the sQTL does not affect the isoform pairs that could potentially undergo NMD or the sum of all weights of the isoform pairs corresponding to the colocalized sQTL, which are identified as candidates for NMD (**Table S3**).

Next, we were interested in protein truncations. For all the 'lacking'-'containing' isoform pairs, we computed the average change in the length of the amino acid sequence between two transcript isoforms or simply the difference in the length of the protein. The resulting pairs of delta length and weight values are then used to compute the weighted average of the delta length, which is stored in the 'Average Delta Amino Acid' field. If there are no delta length and weight pairs, then the value is set to 0 (**Table S4**).

### Supplemental note 10: Experimental validation of *TPM2* in hFOBs

We observed 7 colocalizing sQTLs in total, 5 of which are around exons 5-8 observed in 13-33 tissues depending on the junction. The other two are around exons 9-11 observed specifically in the brain and testis. In order to ensure that all isoforms of *TPM2* are found, regardless of filtering criteria, we decided to cluster all the isoforms based on each time point and create a relative abundance percentage rather than absolute abundance. We were able to capture all four isoforms, albeit, at a low level of expression in certain time points.

In order to confirm that these isoforms are being translated. We were able to identify 12 exon-mapping TPM2 peptides from our total list of 22 peptide targets (**Table S7**). Of those, 4 are unique to exon 6, 4 unique to exon 7, 2 unique to exon 10, and 2 unique to exon 11. Our proteomics results suggest that all four isoforms of *TPM2* identified are being translated and concurs with expression data suggesting a decrease of *TPM2* abundance as hFOBs mature. The ratios of exon 6 and exon 7 in our proteomics analysis suggest an equal presence, which is not consistent with the transcript abundances reported in our long-read data. These results highlight previously reported discordance in transcript-protein correlations [12]. Taken together, we speculate that the ratios of isoforms containing exon 6 or exon 7 are associated with changes in BMD.

### Supplemental methods

### Functional annotation of sQTLs and enrichment regulatory regions

BioMart [13] was used to obtain the genomic positions of lead sQTLs associated with protein-coding genes using GRCh38. Ensembl REST API (https://rest.ensembl.org/) was used to obtain the variants in LD with the lead sQTLs ($r^2 \geq 0.80$; high LD). This information was used as input to GenomicRanges [14] package in R used to identify overlaps of the lead sQTL or those in proxy within the intron (junction) they regulate. We constructed introns from the SQANTI3 GTF file using the package "gread" followed by overlap with splice site acceptor (ssa) or splice site donor (ssd) using GenomicRanges. We used SNPsnap [15] within the package "VSEA" in R to obtain a background set of SNPs with similar minor allele frequencies, distance to nearest genes, and LD patterns. To test enrichment of lead sQTLs splice factor binding sites, we used "fisher.test" within R with a significance threshold alpha < 0.05). The splice factor information was obtained from Van Nostrand et al. [16] and splice factor binding sites were obtained from the eCLIP database within ENCODE [7].
